# Mono-ADP-ribosylation-driven immunosuppression and cross-resistance to therapy through cancer cell intrinsic and extrinsic mechanisms

**DOI:** 10.64898/2026.06.01.729331

**Authors:** Yuchen Sun, Yingying Tang, Vivek T. Singh, Ágnes Holczbauer, Raghavendra Basavaraja, Quoc Thang Bui, Jeong-Hwa Lee, Runnan Gao, A. Cole Edwards, Wei Guo, J. Alan Diehl, Yi Fan, Constantinos Koumenis, Timour Baslan, Ben Z. Stanger, Michael S. Cohen, Vladimir S. Spiegelman, Serge Y. Fuchs

## Abstract

Mono-ADP-ribosylation (MARylation) is emerging as an important regulator of anti-cancer immunity and immunosuppressive tumor microenvironment (TME). Our previous studies showed that PARP11, one of several enzymes that facilitate MARylation, regulates the activities of intratumoral cytotoxic T lymphocytes (CTLs) and regulatory T cells (Tregs). Here, we demonstrate that stimuli such as adenosine, epinephrine, or glucagon-like peptide-1 (GLP1) induced PARP11 in cancer cells. Upregulation of PARP11 in cancer cells led to PARP11-mediated MARylation, ubiquitination, and accelerated degradation of MHC-I through the autophagy-lysosomal pathway. Induction of PARP11 protected cancer cells from killing by specific CTLs and stimulated tumor growth and progression. Genetic ablation of PARP11 attenuated MHC-I MARylation, ubiquitination, and interaction with autophagy receptors. Pharmacologic inhibition of PARP11 in pancreatic ductal adenocarcinoma (PDAC) cells restored their MHC-I levels, sensitized them to killing by CTLs, inhibited tumor growth, and impeded their initial resistance to chemotherapy and their acquired resistance to targeted therapy with RAS inhibitors. Moreover, inhibition of PARP11 prevented hyperprogressive disease in a mouse melanoma model treated with immune checkpoint inhibitors (ICBs), suggesting that PARP11 is a major therapeutically actionable driver of immunosuppression in tumors.

**SYNOPSIS:** Induction of PARP11 in the tumor microenvironment mediates immunosuppression. This study reports that PARP11-driven MARylation and ubiquitination of MHC-I in cancer cells drives immune evasion, tumor growth and resistance to therapies.

## INTRODUCTION

Tumors employ diverse evasive strategies to escape immune surveillance by cytotoxic T lymphocytes (CTLs) in the tumor microenvironment (TME) (1–3). These strategies are focused on suppressing presentation of tumor antigens to undermine the ability of CTLs to recognize cancer cells (1), as well as on targeting CTLs recruitment and their function through CTL exclusion, metabolic inactivation and exhaustion (3, 4). These mechanisms drive tumor resistance to immune therapies as well as to other anti-cancer regimens including chemotherapies, radiotherapies and targeted therapies that also rely on functional anti-tumor CTLs (4–6). Overcoming this critically important problem requires identifying and characterizing key suppressors of CTL-mediated control of tumor growth.

Mono-ADP-ribosyl transferases such as PARP7, PARP14, and PARP11 have recently emerged as important and actionable regulators of the immunosuppressive TME (7). Small molecule inhibitors of PARP7 restore the ability of cancer cells to produce type I interferons (IFN) (8) whereas inactivation of PARP14 prevents resistance to immune checkpoint blockade (ICB) in response to chronic IFN-γ signaling (9). PARP11 drives NAD^+^-dependent mono-ADP ribosylation (MARylation) of diverse substrates, such as the type I IFN receptor IFNAR1 (10), leading to the inactivation of intratumoral CTLs (11) and supporting the suppressive function of Treg cells (12). Genetic or pharmacologic inhibition of PARP11 in T cells disrupts immunosuppressive TME and improves responses to cancer immunotherapies, including ICB and CAR T cells (11, 12).

The selective inhibitor of PARP11, ITK-7 (13), is well tolerated (12) and acts on tumor-infiltrating T cells to increase IFN-γ production (11, 12), stimulate CTLs (14) and inactivate Treg cells (15, 16). However, translating PARP11 inhibitors into the clinic requires a more holistic understanding of how they work, including whether their therapeutic effects extend beyond T cells to include direct actions in malignant cells. A significant group of melanoma and lung cancer patients undergoing ICB therapy experience hyper-progressive disease (HPD, (17)), linked to unique features of cancer cell signal transduction cross-wiring that allow IFN-γ, produced by reactivated T cells, to promote malignant cell growth (18). These mechanisms rely on the depletion of NAD^+^ levels in malignant cells (18). Given that NAD^+^ is an essential substrate of PARP11 (19, 20) and PARP11 inhibitors stimulate IFN-γ production by tumor-infiltrating T cells (11, 12), it is of critical importance to investigate how PARP11 acts in cancer cells.

Here we demonstrate that PARP11 is highly expressed in colon and pancreatic adenocarcinoma cells compared to their normal tissue counterparts. PARP11 is induced in malignant cells by immunosuppressive (adenosine) or stress (epinephrine) mediators, as well as the widely used medication GLP1. Induction of PARP11 in cancer cells enables them to escape CTL surveillance by promoting MARylation, ubiquitination, and lysosomal breakdown of MHC-I proteins. Conversely, PARP11 inhibition prevents the loss of MHC-I and predisposes cancer cells to killing by specific T cells. PARP11 inhibitor restores MHC-I levels in pancreatic ductal adenocarcinoma (PDAC) cells and prevents the development of therapeutic cross-resistance to chemotherapy and targeted therapy in PDAC tumors. Lastly, pharmacologic or genetic inhibition of PARP11 prevents ICB-induced acceleration of tumor growth in a mouse model of HPD.

## RESULTS

### Induction of PARP11 in malignant cells promotes tumor growth in immunocompetent mouse models

High levels of PARP11 expression are associated with poor prognosis in patients with bladder cancer and melanoma (12) and in some other cancer types (21). Increased expression of PARP11 in many types of tumors relative to corresponding normal tissues has also been previously reported (21). Accordingly, in mouse models, we observed higher *Parp11* expression in the colon adenocarcinoma cell lines CT26 and MC38 compared to normal mouse colon (**Fig 1A**). Likewise, murine PDAC cells including 4662, MH6499c4 and MH6419c5 harbored a greater *Parp11* mRNA level compared to normal pancreas (**Fig 1B**).

**Figure 1.**
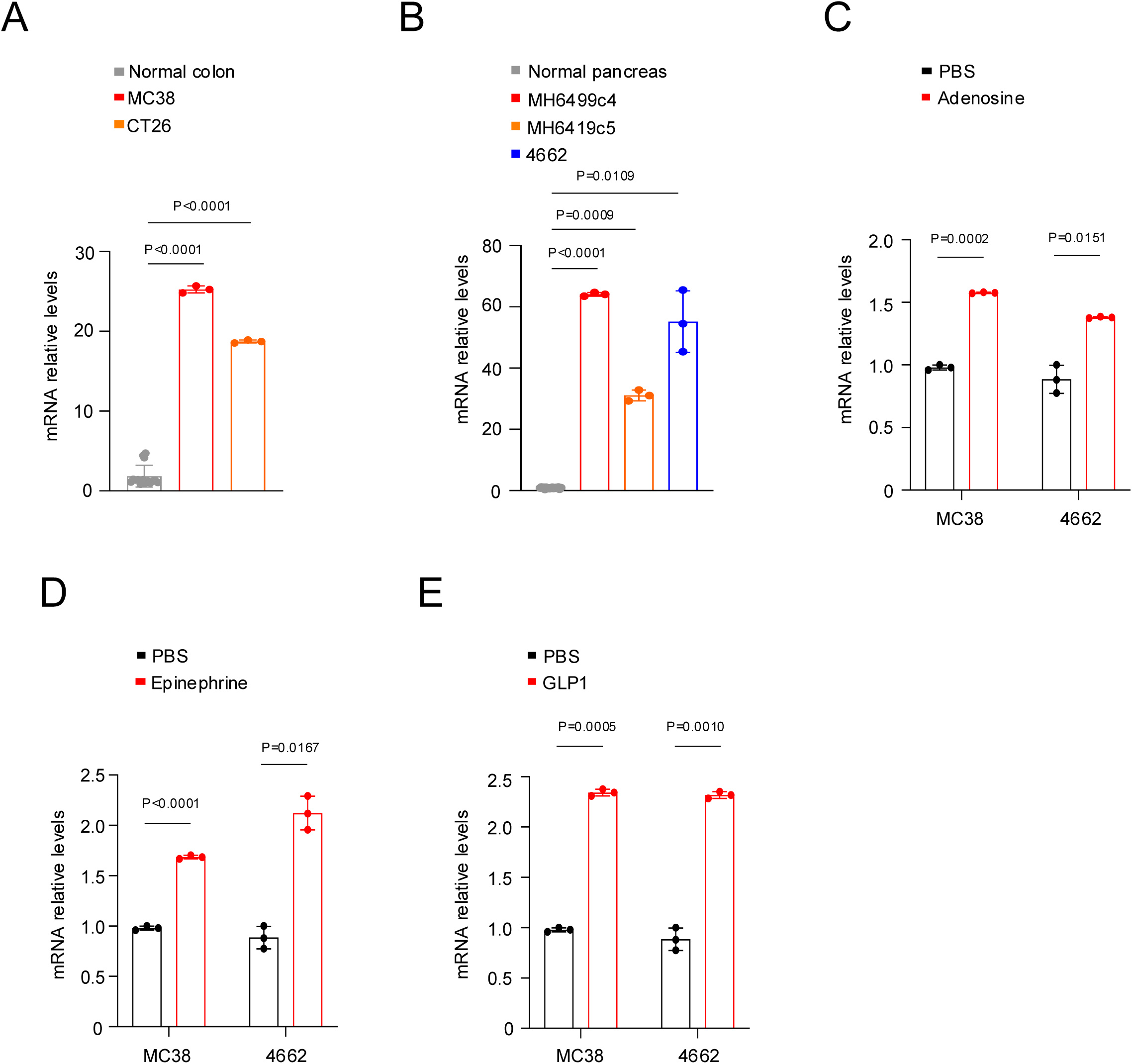
Diverse stimuli in the TME induce PARP11 expression in malignant cells. (A) qRT-PCR analysis of *Parp11* mRNA expression in colon adenocarcinoma cell lines (CT26 and MC38) (n=3) and normal mouse colon tissues (n=15). Data are presented as fold change relative to the normal colon tissues group. (B) qRT-PCR analysis of *Parp11* mRNA expression in pancreatic ductal adenocarcinoma cell lines (4662, MH6499c4, and MH6419c5) (n=3) and normal mouse pancreas tissues (n=18). Data are presented as fold change relative to the normal pancreas group. (C) qRT-PCR analysis of *Parp11* mRNA levels in MC38 and 4662 cells following treatment with adenosine (100 μM for 4 h). Each group contained three biological replicates. Data are presented as fold change relative to the vehicle group. (D) qRT-PCR analysis of *Parp11* mRNA levels in MC38 and 4662 cells following treatment with epinephrine (10 μM for 4 h). Each group contained three biological replicates. Data are presented as fold change relative to the vehicle group. (E) qRT-PCR analysis of *Parp11* mRNA levels in MC38 and 4662 cells following treatment with GLP-1 (7–37) (100 nM for 4 h). Each group contained three biological replicates. Data are presented as fold change relative to the vehicle group. Relative mRNA levels were normalized to Gapdh. Each dot represents one biological replicate. Data are presented as mean ± SEM (A–E). Statistical significance was determined using two-tailed unpaired Student’s t test (A–E).

We next sought to identify stimuli that can induce PARP11 in malignant cells in the TME. The immunosuppressive mediators capable of activating the G-protein coupled receptors (GPRC) and protein kinase A, such as adenosine and prostaglandin E2, upregulated PARP11 in the Treg cells (12). We observed a similar induction of *Parp11* mRNA in MC38, and 4662 malignant cells treated with adenosine (**Fig 1C**). We also investigated the effects of additional physiologically or pharmacologically relevant GPRC activators. Given that adrenergic stress is implicated in immunosuppression and tumor growth and progression (22, 23), we tested the effects of catecholamine treatment. Treatment of MC38 and 4662 cells with epinephrine significantly upregulated *Parp11* expression (**Fig 1D**).

As more than 35% of all approved drugs target GPCR (24, 25), we considered a possibility that some of these agents can modulate PARP11 expression in cancer cells. We selected agents such as glucagon-like peptide-1 (GLP1), which is widely used to treat type 2 diabetes and obesity (26). GLP1 significantly upregulated *Parp11* mRNA in MC38 and 4662 cells (**Fig 1E**). Taken together, these data indicate that diverse stimuli can increase PARP11 levels and, plausibly, function in malignant cells within the TME.

To interrogate the importance of PARP11 in cancer cells, we have generated a battery of isogenic MC38- and CT26-based cell lines in which we either knocked out PARP11 (**Fig S1A-B**) or overexpressed wild-type PARP11 (WT) or its ADP ribosylation-deficient mutant (H197P/Y229S, HY) (**Fig S1C-D**). No notable differences in proliferation rates (**Fig S1E-F**), migration, or invasion through Matrigel in the Boyden chambers (**Fig S1G-H**) were observed between these cell lines.

However, knockout of PARP11 in MC38 cells significantly suppressed their ability to form subcutaneous tumors in the syngeneic C57BL6 mice (**Fig 2A**, **S2A**). Furthermore, similar results were obtained in tumors formed by CT26 cells in the syngeneic Balb/c mice (**Fig 2B**, **S2B**). These observations suggest that PARP11 expression in malignant cells is essential for efficient tumorigenesis.

**Figure 2.**
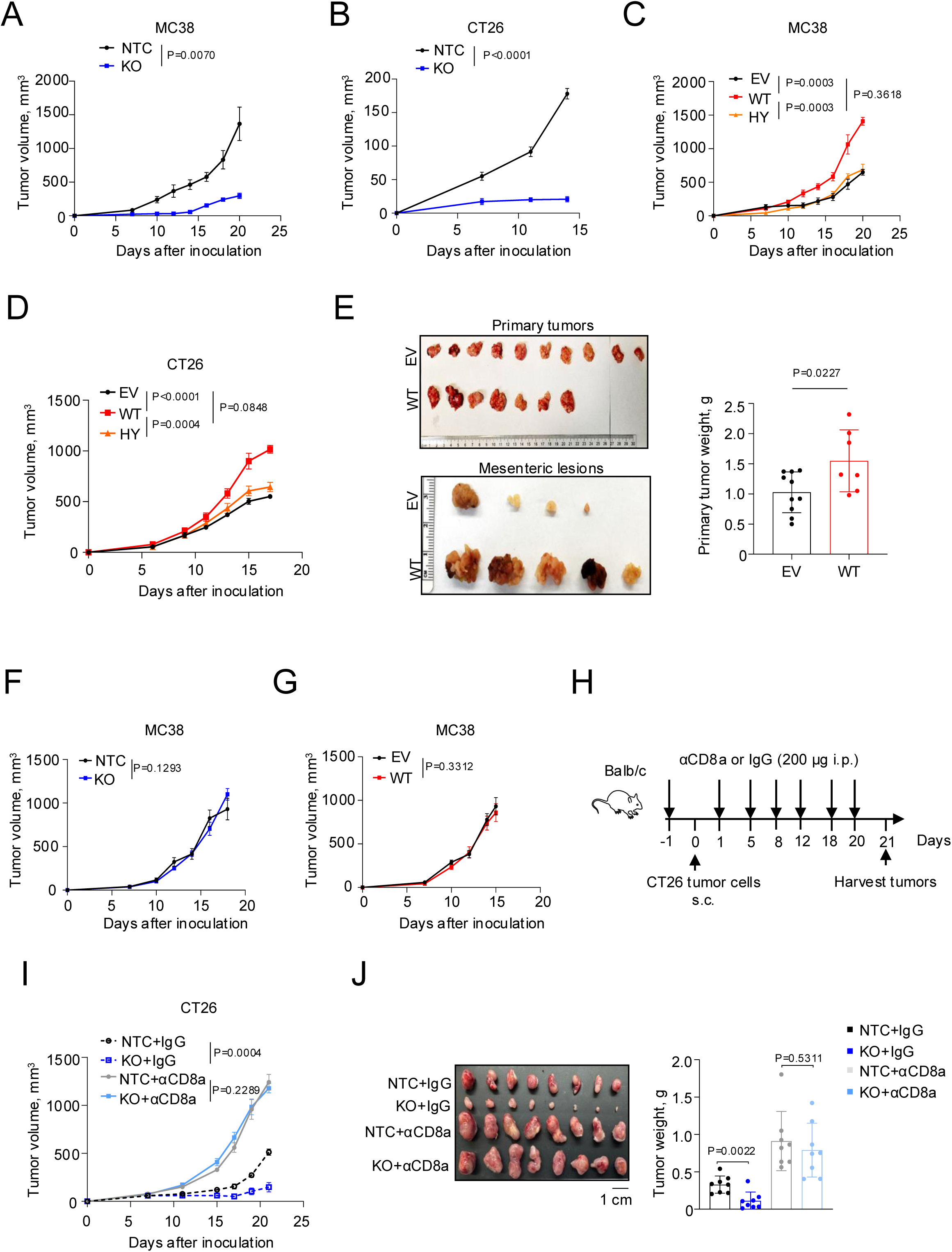
PARP11 promotes tumor growth by evading CD8^+^ T cell-dependent anti-tumor immunity in the TME. (A) Growth curves of subcutaneous MC38 tumors with SgParp11 or NTC guides inoculated into C57BL6 mice (n = 5 per group). (B) Growth curves of subcutaneous CT26 tumors with SgParp11 or NTC guides inoculated into Balb/c mice (n = 5 per group). (C) Growth curves of subcutaneous MC38 tumors overexpressing wild-type PARP11, PARP11 HY mutant, or EV control inoculated into C57BL6 mice (n = 5 per group). (D) Growth curves of subcutaneous CT26 tumors overexpressing wild-type PARP11, PARP11 HY mutant, or EV control into Balb/c mice (n = 7 per group). (E) Representative images and weighs of primary tumors and images of mesenteric metastatic lesions in Balb/c mice orthotopically inoculated with CT26 tumor cells overexpressing wild-type PARP11 (n = 7) or EV control (n = 10). (F) Growth curves of subcutaneous MC38 tumors with SgParp11 or NTC guides inoculated into *Rag1*^−/−^ mice (n = 7 per group). (G) Growth curves of subcutaneous MC38 tumors overexpressing wild-type PARP11 or EV control inoculated into *Rag1*^−/−^ mice (n = 7 per group). (H) Schematic illustration of the treatment schedule for IgG control or anti-CD8α antibody treatment in Balb/c mice bearing subcutaneous CT26 tumors with SgParp11 or NTC guides (n = 8 per group). (I) Growth curves of subcutaneous CT26 tumors with SgParp11 or NTC in Balb/c mice treated with either IgG control or anti-CD8α antibody as described in (H). (J) Representative tumor images and tumor weights of subcutaneous CT26 tumors with SgParp11 or NTC in mice treated as described in (H). Each dot represents one biological replicate. Data are presented as mean ± SEM. Statistical significance was determined by ANOVA, followed by Tukey’s multiple comparisons (A-D, F, G, I). Statistical significance was determined using two-tailed unpaired Student’s t test (E) and (J).

Consistent with this hypothesis, overexpression of WT PARP11 in MC38 cells significantly accelerated subcutaneous tumor growth in syngeneic C57BL6 mice (**Fig 2C**, **S2C**). Similar results were observed for CT26 cells in the syngeneic Balb/c mice (**Fig 2D**, **S2D**). Furthermore, PARP11-overexpressing CT26 cells also formed larger primary tumors and produced more and larger local metastatic mesenteric lesions when inoculated orthotopically into the ileocecal corner (**Fig 2E**). In both MC38 and CT26 models, the MARylation-deficient PARP11 HY mutant was expressed at levels similar to WT (**Fig S1C-D**) and did not accelerate tumor growth (**Fig 2C-D, S2C-D**). These results suggest that induction of PARP11 in cancer cells stimulates tumorigenesis and that the MARylation function of PARP11 is essential for supporting this phenotype.

Importantly, neither knockout (**Fig 2F, S2E**) nor overexpression (**Fig 2G, S2F**) of PARP11 significantly altered tumor growth of MC38 cells inoculated into the lymphocyte-deficient *Rag1*^-/-^ mice. These results argue for the importance of adaptive immunity in the tumorigenesis driven by the induction of PARP11 in cancer cells. To test this possibility in another model, we depleted CD8^+^ T cells from Balb/c mice bearing CT26 tumors (**Fig 2H**). This depletion did not affect the levels of natural killer (NK) cells or CD4^+^ T cells in peripheral blood or spleens of these mice (**Fig S2G**) yet sufficed to accelerate growth of CT26 tumors regardless of knockout of PARP11 (**Fig 2I-J**). Overall, these results indicate that induction of PARP11-driven MARylation in malignant cells stimulates tumor growth by evading anti-tumor CD8-dependent immune responses.

### Induction of PARP11 in malignant cells aids in downregulating MHC-I and evading anti-tumor CTLs

PARP11-driven evasion of anti-tumor immunity may arise from the ability of malignant cells to actively repress anti-tumor CD8^+^ T cells or/and to undermine their ability to recognize cancer cells. No significant changes in the numbers of splenic T cells were observed in the spleens of mice bearing tumors from CT26 or MC38 cells harboring PARP11 overexpression or knockout (**Fig S3A**) arguing against systemic immune suppression as an underlying mechanism of the observed tumor phenotypes.

PARP11-dependent MARylation inhibits the type I interferon pathway (10), which is known to regulate the expression of chemokines (such as CXCL9/10/11 and CCL2/5/7) that can stimulate tumor infiltration with T lymphocytes and myeloid cells (27). We immunoprofiled tumors formed by CT26 or MC38 cells (**Fig S3B**) and found that overexpression of PARP11 did not elicit notable changes in the frequency or numbers of tumor-infiltrating myeloid cells (**Fig S3C**) or CD4^+^ T cells or NK cells (**Fig S3D**). However, tumors formed by CT26 or MC38 cells overexpressing WT but not HY mutant PARP11 displayed a modest yet significant decrease in frequency of CD8^+^ T cells (**Fig 3A**) as well in their expression of Granzyme B or IFN-γ (**Fig 3B**). These results suggest that the numbers and activities of tumor-infiltrating CD8^+^ CTLs can be attenuated by PARP11-driven MARylation in cancer cells.

**Figure 3.**
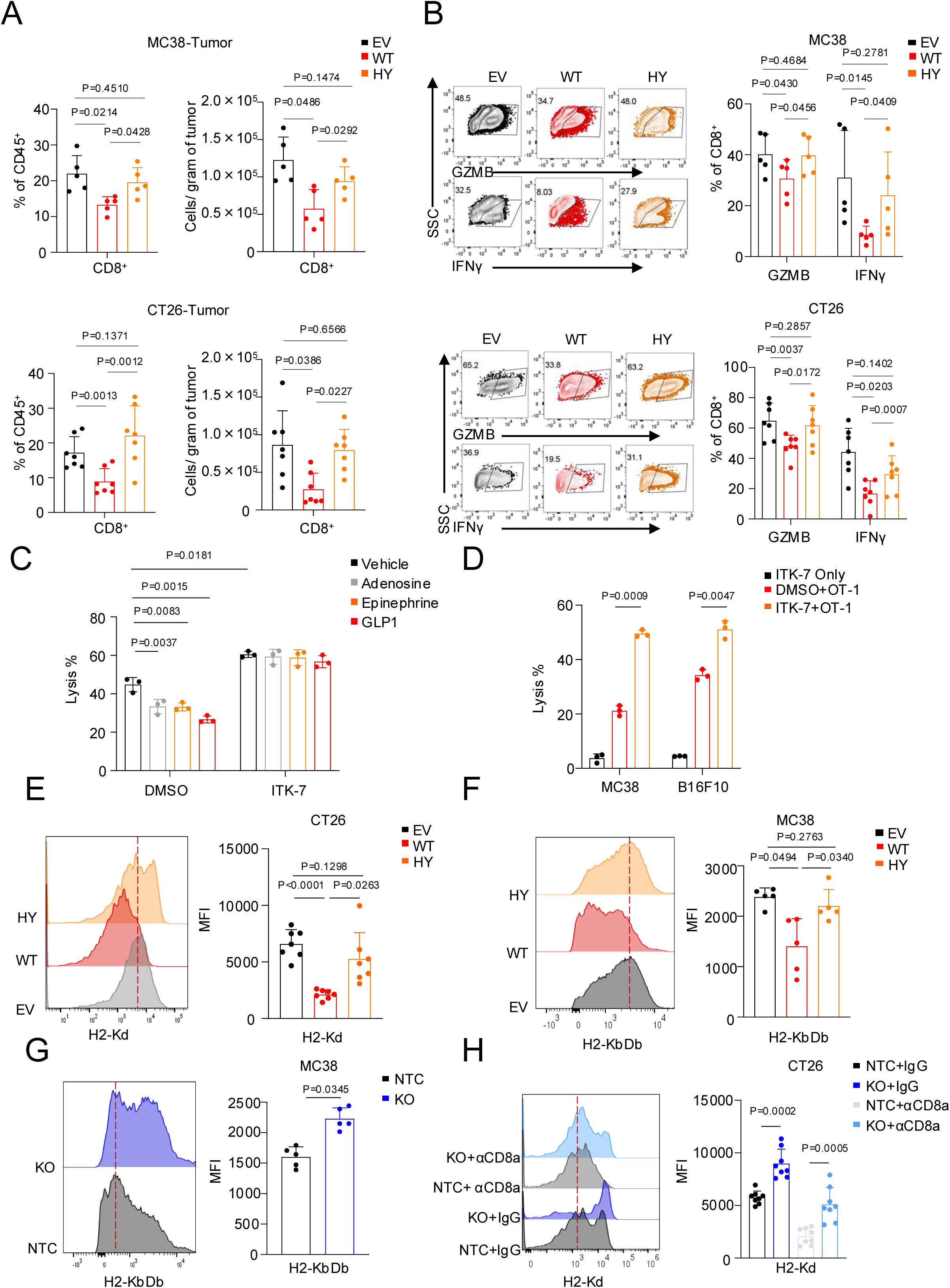
PARP11 suppresses malignant cell surface MHC-I and promotes CD8^+^ T cell-mediated immune evasion. (A) Flow cytometry analysis of the frequency (% of live CD45^+^ cells) and the number (per gram of tumor tissue) of CD8^+^ T cells from subcutaneous MC38 or CT26 tumors overexpressing wild-type PARP11, PARP11 HY mutant, or EV control in C57BL6 (n = 5) or Balb/c (n = 7) mice, as described in Figure 2C and 2D. (B) Flow cytometry analysis of the percentage of IFN-γ^+^ and Granzyme B^+^ CD8^+^ T cells isolated from subcutaneous MC38 or CT26 tumors overexpressing wild-type PARP11, PARP11 HY mutant, or EV control in C57BL6 (n = 5) or Balb/c (n = 7) mice, as described in Figure 2C and 2D. (C) Killing of MC38-OVA-luc cells pre-treated or not with ITK-7 (1 μM, 48 h) or vehicle, followed by adenosine (100 μM), or epinephrine (10 μM), or GLP-1 (100 nM) before co-culture with OT-1 CTLs for 12 h (OT-1: MC38-OVA-luc = 5:1; n = 3). (D) Killing of MC38-OVA-luc cells or B16F10-OVA-luc cells pretreated with ITK-7 (1 μM, 48 h) or vehicle prior to co-culture with OT-1 CTLs for 12 h (OT-1: MC38-OVA-luc = 5:1; OT-1: B16F10-OVA-luc = 5:1; n = 3). (E) Flow cytometry analysis of mean fluorescence intensity (MFI) of H2-Kd on malignant (CD45^−^ Podoplanin^+^) cells from subcutaneous CT26 tumors overexpressing wild-type PARP11, PARP11 HY mutant, or EV control in Balb/c mice (n = 7), as described in Figure 2D. (F) Flow cytometry analysis of MFI of H2-KbDb on malignant (CD45^−^EpCAM^+^) cells from subcutaneous MC38 tumors overexpressing wild-type PARP11, PARP11 HY mutant, or EV control in C57BL6 mice (n = 5), as described in Figure 2C. (G) Flow cytometry analysis of MFI of H2-KbDb on malignant (CD45^−^EpCAM^+^) cells from subcutaneous MC38 tumors expressing sgParp11 or NTC in C57BL6 mice (n = 5), as described in Figure 2A. (H) Flow cytometry analysis of MFI of H2-Kd on malignant (CD45^−^PDPN^+^) cells from subcutaneous CT26 tumors treated as described in Figure 2H (n = 8). Each dot represents one biological replicate. Data are presented as mean ± SEM. Statistical significance was determined using two-tailed unpaired Student’s t test (A–H).

The presence of tumor-infiltrating CTLs in human cancers positively correlates with MHC-I expression on cancer cells (1, 28). These CTLs recognize tumor-associated antigen peptides presented on the cell surface in complex with β2-microglobulin and MHC-I transmembrane proteins (1). We next used an in vitro killing assay (11, 12) to examine whether PARP11 affects the MHC-I-mediated recognition of ovalbumin-expressing cancer cells by specific cytotoxic OT-1 CTLs. In these settings, pre-treatment of MC38-OVA target cells with stimuli capable of inducing PARP11 (such as adenosine, epinephrine or GLP1) prior to exposing them to OT-1 T cells notably decreased the extent of killing (**Fig 3C**).

OT-1 CTLs exposed to MC38-OVA or B16F10-OVA target cells treated or not with ITK-7 expressed comparable levels of cytotoxic mediators Granzyme B, IFN-γ or perforin (**Fig S3E**). While pre-treatment of target cancer cells with ITK-7 alone did not cause cytotoxicity, it significantly increased their killing by specific OT-1 CTLs (**Fig 3D**). These results collectively suggest that PARP11-driven MARylation aids cancer cells in evading their recognition by CTLs and protects cancer cells from ensuing cytotoxicity.

Given that antigen-specific killing is mediated by the MHC-I complex, we next examined whether PARP11 regulates MHC-I levels. Flow cytometry comparison of splenocytes from naïve WT or *Parp11*^-/-^ mice revealed that the latter expressed more MHC-I on their surface (**Fig S3F**). Conversely, treatment of MC38 cells with PARP11 inducers such as adenosine, epinephrine and GLP1 significantly decreased MHC-I on cell surface (**Fig S3G**) suggesting that these stimuli might co-opt PARP11 in undermining the presentation of tumor antigens by cancer cells.

We next analyzed the in vivo cell-surface levels of the haplotype-specific MHC-I proteins in MC38 (H2-KbDb) and CT26 (H2-Kd) cells isolated from the corresponding tumors. We found that overexpression of WT, but not HY mutant, PARP11 significantly downregulated MHC-I proteins on the surface of PDPN^+^ or EPCAM^+^ malignant cells in CT26 tumors (**Fig 3E**) or in MC38 tumors (**Fig 3F**). Conversely, knockout of PARP11 upregulated cancer cell surface MHC-I in tumors formed by PARP11-knockout MC38 cells (**Fig 3G**). Likewise, PARP11-deficient CT26 cells displayed increased MHC-I in tumors regardless of whether CD8^+^ T cells were depleted in the host mice or not (**Fig 3H**). These results indicate an unexpected role for PARP11 in regulating MHC-I and may explain how PARP11 induction helps cancer cells evade CTLs.

### PARP11 promotes MARylation, ubiquitination and lysosomal degradation of MHC-I

We next sought to delineate the mechanisms through which induced PARP11 can downregulate the cell surface levels of MHC-I. Levels of mRNA for the genes encoding MHC-I, β2-microglobulin or elements of antigen peptide presentation machinery (such as Tap1, Tap2 or Tapbp) were not affected either by over-expression or knockout of PARP11 (**Fig S4A-B**). However, MHC-I levels are subject to post-translational regulation via ubiquitin-driven degradation (reviewed in (28, 29)). Importantly, ADP-ribosylation emerged as a key regulator of the ubiquitination-dependent proteolysis (30). Thus, we focused on post-translational regulation of MHC-I.

We found that overexpression of WT PARP11 in CT26 or MC38 cells decreased total MHC-I protein levels, whereas the HY mutant had a notably smaller effect (**Fig 4A**). Evaluation of the half-life of MHC-I protein in MC38 cells using cycloheximide chase revealed that overexpression of WT PARP11 significantly accelerated MHC-I proteolytic turnover. Importantly, this effect was attenuated by treatment with PARP11 inhibitor ITK-7 (**Fig 4B**), suggesting that PARP11-driven MARylation promotes the degradation of MHC-I protein.

**Figure 4.**
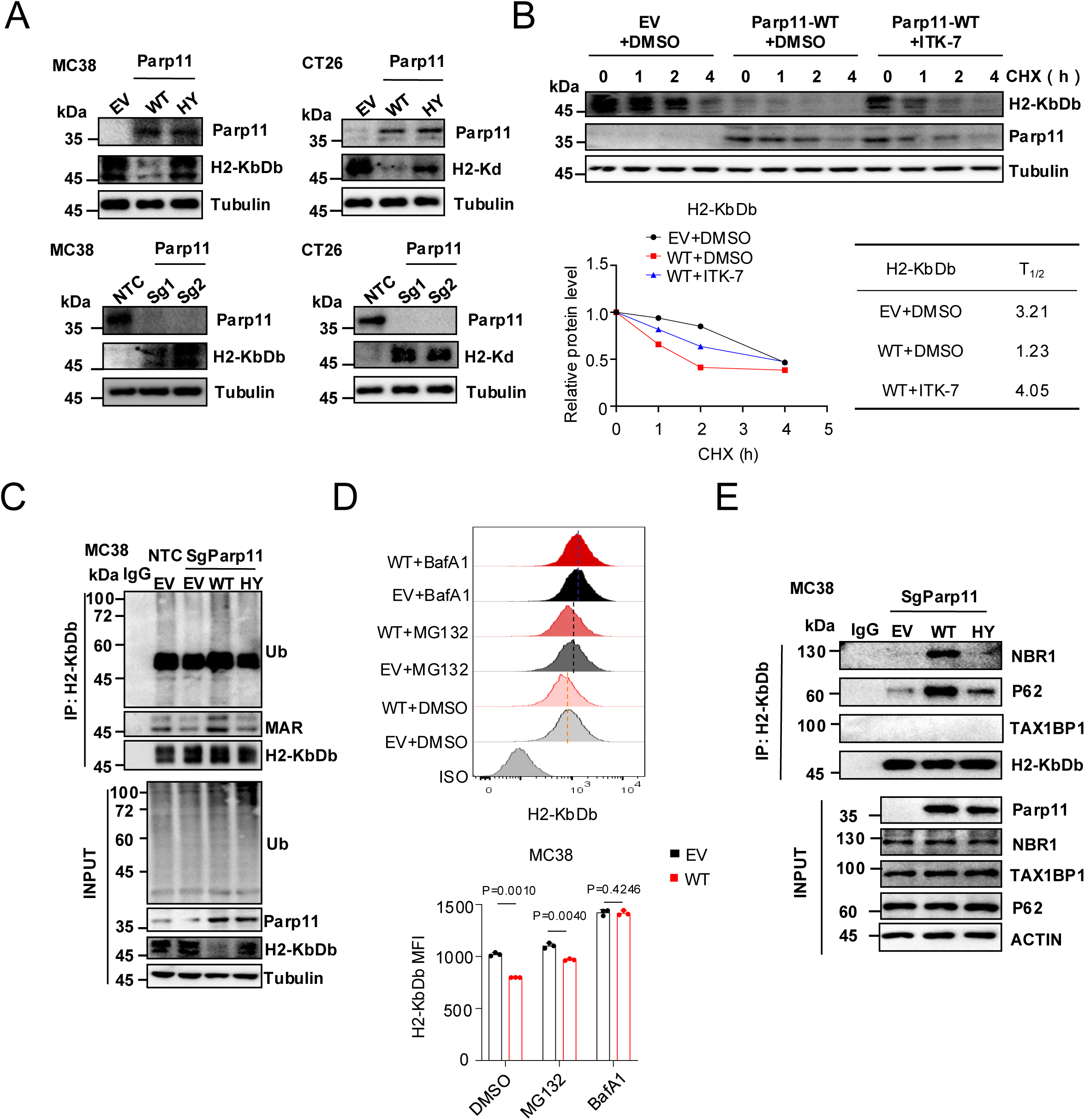
PARP11 promotes MHC-I ubiquitination and lysosomal degradation through MARylation. (A) Immunoblot analysis of H2-KbDb or H2-Kd expression in MC38 and CT26 cells with *Parp11* knockout, wild-type PARP11 overexpression, PARP11 HY mutant overexpression, or corresponding controls, as described in Figure S1A–S1D. α-Tubulin was used as a loading control. (B) Immunoblot analysis of H2-KbDb expression in MC38 cells overexpressing wild-type PARP11 or EV control following treatment with ITK-7 (1 μM 48h) or DMSO and then with cycloheximide (CHX, 50 μg/mL) for the indicated time points. Immunoblot signals were quantified using ImageJ, and the relative H2-KbDb protein levels were plotted over time to determine protein half-life. Data are presented as fold change relative to the control group. (C) MC38 cells with endogenous *Parp11* knockout were reconstituted with wild-type PARP11, PARP11 HY mutant, or EV control were characterized by direct immunoblotting (“Input”, lower panel). Analyses of H2-KbDb MARylation and ubiquitination were carried out on MHC-I immunoprecipitates equilibrated for H2-KbDb levels (upper panel). (D) Flow cytometry analysis of MFI of H2-KbDb on MC38 cells overexpressing wild-type PARP11 or EV control pretreated with MG132 (5 μM), bafilomycin A1 (BafA1, 100 nM), or vehicle for 4 h (n = 3). (E) Immunoprecipitation-immunoblotting analyses of interaction between H2-KbDb and NBR1, p62 or TAXBP1 autophagy acceptors in MC38 cells with endogenous Parp11 knockout were reconstituted with wild-type PARP11, PARP11 HY mutant, or EV control. Direct immunoblotting control (“Input”, lower panel) is also shown. Each dot represents one biological replicate. Data are presented as mean ± SEM. Statistical significance was determined using two-tailed unpaired Student’s t test for (D).

Along with genetic defects and transcriptional regulation, the accelerated ubiquitination-driven lysosomal or proteasomal degradation of MHC-I is a common mechanism through which cancer cells evade anti-tumor immunity, sustain their growth and resist immune therapies (reviewed in (2, 28, 29, 31)). The interplay between protein ADP-ribosylation and ubiquitination has emerged as an important regulator of diverse biological processes (32). To evaluate the importance of PARP11 in MARylation and ubiquitination, we generated a battery of four isogenic MC38- or CT26-based cell lines that included control cells, PARP11 knockout cells and the latter engineered to express WT or HY PARP11 (**Fig S4C**). In line with results seen in **Fig 4A**, immunoblot analysis of cell lysates revealed that expression of WT PARP11 but not of catalytically inactive HY PARP11 mutant decreased total MHC-I levels (“Input” panels in **Fig 4C** and **S4D**).

We immunopurified MHC-I proteins from these cells, equilibrated to yield equal levels of these proteins loaded on the SDS-PAGE gels and subjected these gels to immunoblot analyses using anti-mono-ADP-ribose and anti-ubiquitin antibodies. This analysis revealed that PARP11 knockout decreased MARylation and ubiquitination of MHC-I. This phenotype was reversed by expression of WT but not HY PARP11 in MC38 (**Fig 4C**) or in CT26 (**S4D**) cells. In all, these data suggest that PARP11 controls MARylation and ubiquitination of MHC-I.

In many cancers, the loss of MHC-I occurs through a lysosomal pathway. In these cells, ubiquitinated MHC-I is recognized and trafficked to lysosomes for degradation by ubiquitin-binding autophagy cargo receptors (33). We observed that downregulation of MHC-I in PARP11-expressing MC38 (**Fig 4D**) or CT26 (**Fig S4E**) cells was reversed by treatment with the lysosomal inhibitor Bafilomycin A1, but not by the proteasomal inhibitor MG132. Accordingly, we used a co-immunoprecipitation analysis to assess the recruitment of autophagy receptors, including NBR1, p62, and TAX1BP1, to MHC-I (**Fig 4E**). We did not detect TAX1BP1 in MHC-I immunoprecipitates; however, genetic studies in isogenic PARP11-deficient MC38 (**Fig 4E**) or CT26 (**Fig S4F**) cells revealed that re-expression of WT (but not HY) PARP11 increased the interactions of MHC-I with p62 or/and NBR1 autophagy cargo receptors.

In all, these data suggest that PARP11-driven MARylation stimulates MHC-I ubiquitination and drives its subsequent lysosomal degradation. It is plausible that the resulting MHC-I downregulation could render these cells less visible to specific CTLs, thereby helping them evade anti-tumor immune responses and stimulate tumor growth. To test this hypothesis, we next expressed exogenous H2-Kb and β2-microglobulin proteins in the MC38-OVA isogenic cell lines to equalize MHC-I cell-surface levels (**Fig. S5A**).

No difference in cell death was noted in these cells cultured alone (**Fig 5A**). However, when these cells were co-cultured with OT-1 CTLs, we observed that PARP11 knockout predisposed target cells for killing whereas re-expression of PARP11 attenuated the cytotoxic effects (**Fig 5A**). Importantly, these phenotypes were abolished in cell lines with equalized cell-surface MHC-I levels (**Fig 5A**). These results suggest that PARP11-driven MARylation helps cancer cells evade the CTLs in a manner that at least partially depends on downregulation of MHC-I.

**Figure 5.**
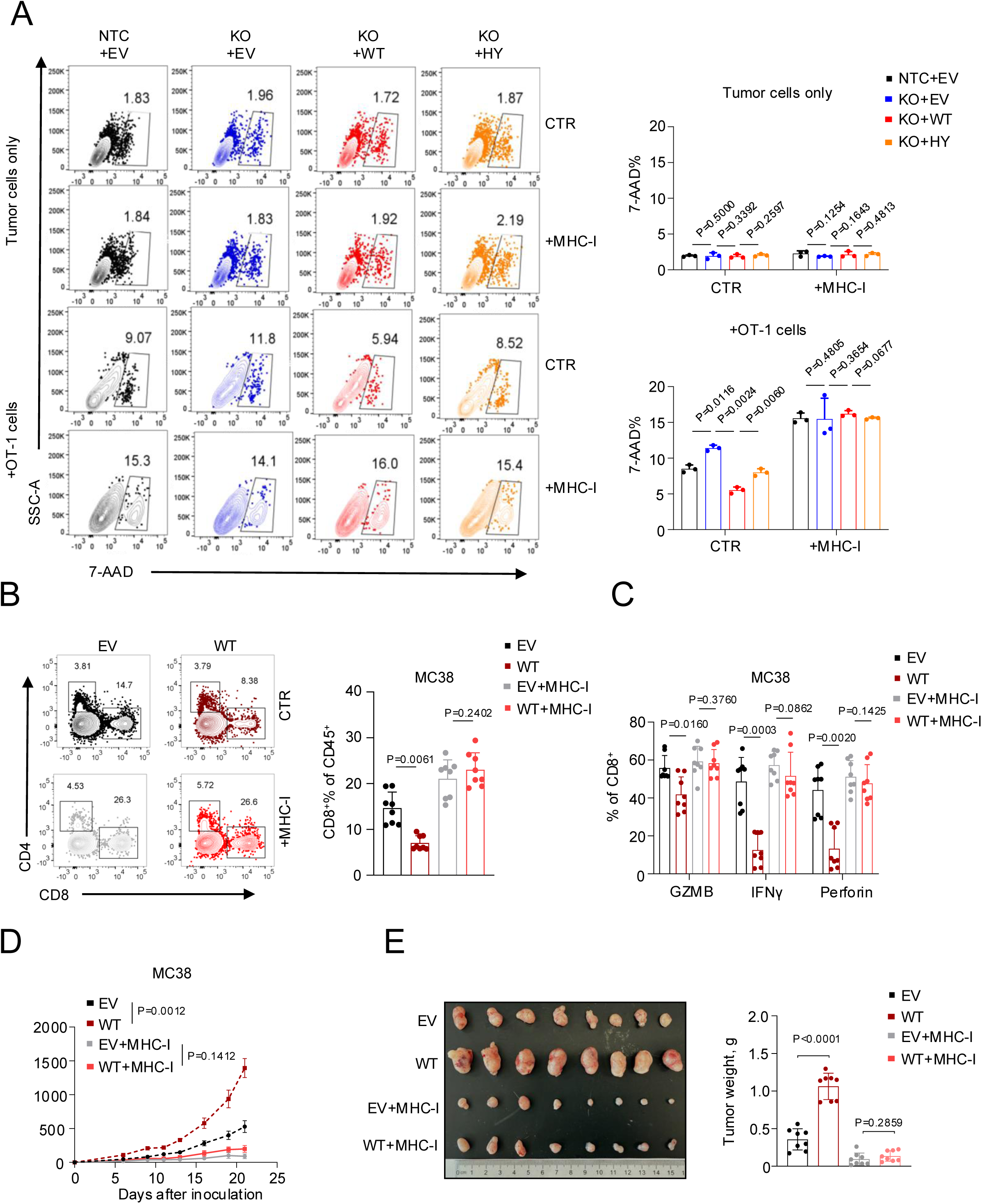
Restoration of MHC-I reverses PARP11-mediated immune evasion. (A) MC38-OVA cells with endogenous *Parp11* knockout were reconstituted with wild-type PARP11, PARP11 HY mutant, or EV control, followed by overexpression of H2-Kb and β2-microglobulin to restore equivalent MHC-I surface levels. Killing of these cells by co-cultured OT-1 CTLs (12 h, OT-1: MC38-OVA = 1:1) was assessed by flow cytometry analysis of the frequency of dying 7-AAD^+^ MC38-OVA cells (n = 3 per group). (B) Flow cytometry analysis of the frequency (% of live CD45^+^ cells) of CD8^+^ T cells in subcutaneous MC38 tumors in C57BL6 mice (n = 8). MC38 cells overexpressing wild-type PARP11 or EV control were further transduced with H2-Kb and β2-microglobulin to equalize tumor cell surface MHC-I levels. (C) Flow cytometry analysis of the percentage of Granzyme B^+^, IFN-γ^+^, and Perforin^+^ CD8^+^ T cells isolated from subcutaneous MC38 tumors in C57BL6 mice treated as described in (B). (D) Tumor growth curves of subcutaneous MC38 tumors in C57BL6 mice (n = 8 per group) treated as described in (B). (E) Representative tumor images and tumor weights of subcutaneous MC38 tumors from C57BL6 mice (n = 8 per group) treated as described in (B). Each dot represents one biological replicate. Data are presented as mean ± SEM. Statistical significance was determined using two-tailed unpaired Student’s t test for (A–C) and (E), and one-way ANOVA followed by Tukey’s multiple-comparisons test for (D).

To test this possibility in vivo, we compared the CTLs infiltration in MC38 tumors re-expressing PARP11 with or without co-expression of MHC-I. Flow cytometry analysis revealed that co-expression of MHC-I attenuated the decrease in CD8^+^ T cell infiltration in tumors expressing PARP11 (**Fig 5B** and **S5B**). Furthermore, analysis of cytotoxic markers revealed that expression of PARP11 in malignant cells decreased the percentage of the intratumoral CD8^+^ CTLs positive for Granzyme B, perforin or IFN-γ. Importantly, these differences were no longer evident in CTLs from tumors formed by cells re-expressing MHC-I (**Fig 5C** and **S5C**). Lastly, re-expression of MHC-I inhibited growth of MC38 tumors regardless of PARP11 expression, which otherwise significantly accelerated tumor growth (**Fig 5D-E**). Overall, these results indicate that PARP11-dependent MARylation controls cell-surface MHC-I and CTL evasion, and that this mechanism accelerates tumor growth.

### Inhibition of PARP11 overcomes cross-resistance of PDAC tumors to therapies and prevents hyper-progressive disease in melanoma model

The highly potent and selective PARP11 inhibitor, ITK-7 (13), was well tolerated and effective at decelerating tumor growth as a single agent or in combination with immune checkpoint blockade (12). While these effects were attributed to PARP11 modulation in T cells, our current data suggest that effects on MHC-I levels in cancer cells may be mechanistically important. Indeed, in vitro pre-treatment of MC38 cells with ITK-7 upregulated cell surface MHC-I and prevented its loss in response to PARP11 inducers such as adenosine, epinephrine, and GLP1 (**Fig S6A**).

Given that accelerated lysosomal degradation of MHC-I is a major mechanism of its inactivation in PDAC tumors (33, 34) and that PARP11 levels were increased in mouse (**Fig 1B**) and human (**Fig S6B**) tumors compared to normal tissue controls, we examined the effects of PARP11 inhibition in these cells. Treatment of mouse PDAC cells with ITK-7 consistently upregulated MHC-I (**Fig S6C**). Similar results were obtained in PANC1, MP2, and HPAC human pancreatic cell lines (**Fig 6A**), suggesting that ITK-7 could be used to pharmacologically restore MHC-I levels in pancreatic cancer cells.

**Figure 6.**
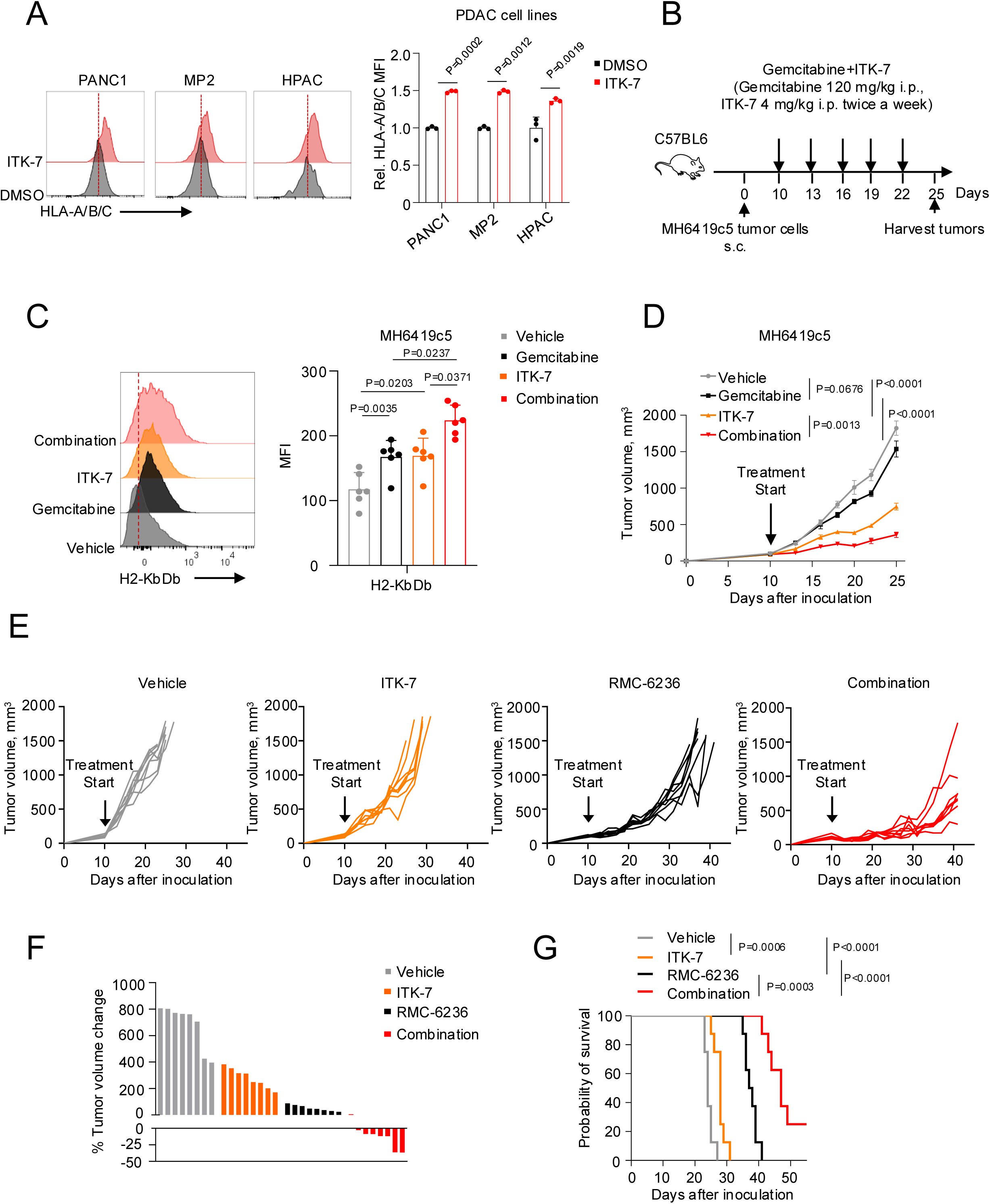
PARP11 inhibition overcomes therapeutic cross-resistance in PDAC tumors. (A) Flow cytometry analysis of MFI of HLA-A/B/C on PANC1, MP2, and HPAC cells pretreated with ITK-7 (1 μM, 48 h) or DMSO. Data are presented as fold change relative to the control group. (B) Schematic illustration of the treatment schedule for administration (intraperitoneally, twice weekly) of gemcitabine (120 mg/kg) and ITK-7 (4 mg/kg) in C57BL6 mice bearing subcutaneous MH6419c5 tumors (n = 6 per group). (C) Flow cytometry analysis of MFI of H2-KbDb on malignant CD45^−^YFP^+^ cells isolated from subcutaneous MH6419c5 tumors treated as described in (B). (D) Tumor growth curves of subcutaneous MH6419c5 tumors treated as described in (B). (E) Individual tumor growth curves of subcutaneous MH6419c5 tumors treated with RMC-6236 (25 mg/kg, oral gavage daily) and ITK-7 (4 mg/kg, intraperitoneally, twice weekly) (n = 8 per group). (F) Waterfall plot showing the percentage change in tumor volume from baseline at the first day of treatment (Day 10) to Day 17 after tumor inoculation. Mice were treated as described in (E). (G) Kaplan-Meier analysis of survival of mice treated as described in (E). Each dot represents one biological replicate. Data are presented as mean ± SEM. Statistical significance was determined using two-tailed unpaired Student’s t test for (A) and (C), and one-way ANOVA followed by Tukey’s multiple-comparisons test for (D). (G) is represented using Kaplan-Meier curves.

Inclusion of ITK-7 in the immune checkpoint blockade regimen reversed resistance to this treatment in the PDAC model of the “cold” MH6419c5 tumor (12). As PARP11 acts to mask cancer cells from CTLs, and the latter play an essential role in the success or failure of the majority of anti-cancer regimens (6), including those used to treat pancreatic cancer (35), we sought to investigate the putative roles of PARP11 in cross-resistance of PDAC. We used the MH6419c5 tumor model, shown to be refractory to chemotherapy with gemcitabine (36) and to develop resistance to RAS inhibitors (37). Treatment of MH6419c5 tumors with gemcitabine (**Fig 6B**) led to a significant increase in cancer cell surface MHC-I levels (**Fig 6C**), indicating an on-target effect. Furthermore, while gemcitabine alone did not significantly affect tumor growth, the combination of gemcitabine plus ITK-7 resulted in a notable deceleration of tumor growth compared with ITK-7 monotherapy (**Fig 6D** and **S6D**).

Hyperactivated oncogenic RAS pathway has long been associated with inhibition of MHC-I-dependent antigen presentation (38, 39). Furthermore, T cells play a key role in the extent and duration of PDAC tumors’ responses to RAS inhibitors (37). Consistent with these studies, daily treatment with the RAS inhibitor RMC-6236 in MH6419c5 tumors initially showed robust anti-tumor effects, but tumors eventually resumed growth. The inclusion of ITK-7 in the RMC-6236 produced significantly stronger combination effects than monotherapy with either agent (**Fig 6E**). Importantly, combining ITK-7 with RMC-6236 significantly extended the lifespan of animals bearing MH6419c5 tumors (**Fig 6F, G**). Overall, these data suggest that inhibition of PARP11 can overcome initial or acquired resistance to chemotherapy or targeted therapy in PDAC.

Zou and colleagues established an immune competent model of the IFN-γ-driven HPD using YUMM1.7 melanoma tumors treated with anti-PD-L1 antibody (18). We examined the effects of ITK-7 treatment in these settings. Administration of ITK7 decreased the number of intratumoral myeloid cells (**Fig S7A**) but increased the levels of tumor-infiltrating T cells (**Fig S7B**).

Furthermore, intratumoral CTLs from ITK-7-treated mice displayed greater expression of IFN-γ (**Fig 7A** and **S7C**). These data suggest that inhibition of PARP11 can stimulate IFN-γ production in the YUMM1.7 model, thereby raising a concern that ITK-7 or similar PARP11 inhibitors may exacerbate HPD.

**Figure 7.**
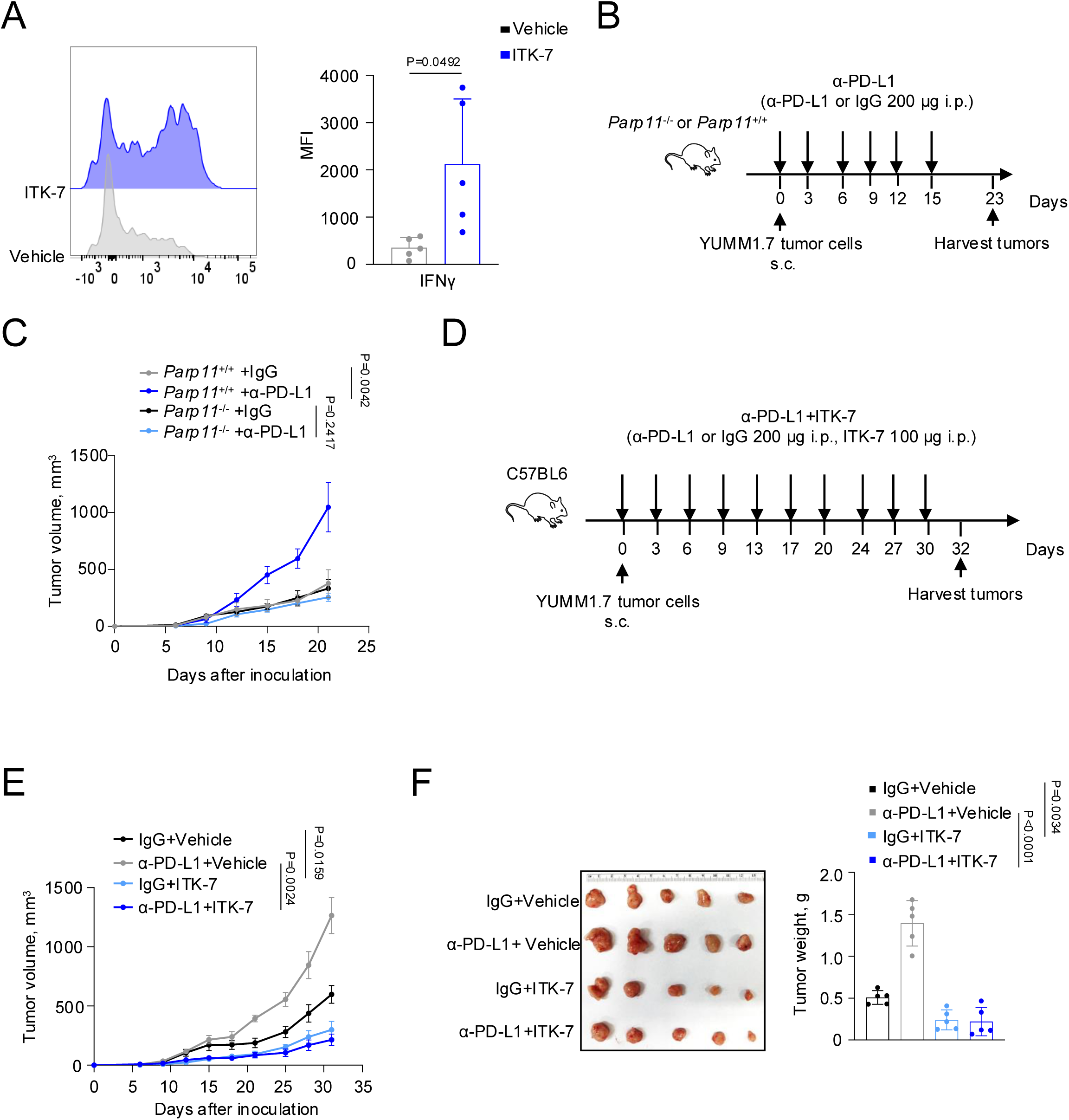
PARP11 inhibition prevents hyper-progressive disease in melanoma. (A) Flow cytometry analysis of MFI of IFN-γ^+^ CD8^+^ T cells isolated from subcutaneous YUMM1.7 tumors grown in WT syngeneic mice treated with anti–PD-L1 antibody (200 μg per mouse, intraperitoneally). Mice were concurrently treated with vehicle control or ITK-7 (100 μg per mouse, intraperitoneally, twice a week for 4 weeks; n = 5 per group). (B) Schematic illustration of the treatment schedule for anti–PD-L1 antibody or IgG control treatment (200 μg per mouse, intraperitoneally) in WT (n = 5) or *Parp11*^−/−^ (n = 4 or 5) mice bearing subcutaneous YUMM1.7 tumors. (C) Tumor growth curves of subcutaneous YUMM1.7 tumors treated as described in (B). (D) Schematic illustration of the treatment schedule for anti–PD-L1 antibody or IgG control (200 μg per mouse, intraperitoneally) together with ITK-7 (100 μg per mouse, intraperitoneally) or vehicle control treatment in C57BL6 mice bearing subcutaneous YUMM1.7 tumors (n = 5 per group). (E) Tumor growth curves of subcutaneous YUMM1.7 tumors treated as described in (D). (F) Representative tumor images and tumor weights of subcutaneous YUMM1.7 tumors from mice treated as described in (D). Each dot represents one biological replicate. Data are presented as mean ± SEM. Statistical significance was determined using two-tailed unpaired Student’s t test for (A) and (F), and one-way ANOVA followed by Tukey’s multiple-comparisons test for (C) and (E).

We tested this hypothesis in subsequent in vivo studies (**Fig 7B**). Consistent with previously published observations (18), treatment of wild-type C57BL6 syngeneic mice bearing YUMM1.7 tumors with an anti-PD-L1 antibody significantly accelerates tumor growth. However, this phenotype was not observed in *Parp11*^-/-^ animals (**Fig 7C** and **S7D**), suggesting that inactivation of PARP11 in the TME does not aggravate HPD. Moreover, given that PARP11-driven MARylation in cancer cells remains unaffected under these conditions, we anticipated an even greater anti-tumor effect of pharmacologic inhibition of PARP11.

Indeed, the accelerated growth of YUMM1.7 tumors in ICB-treated animals (**Fig 7D**) was not further augmented by ITK7 treatment. Moreover, animals that received ITK-7 significantly decelerated tumor growth regardless of whether they were treated with anti-PD-L1 or control antibody (**Fig 7E-F**). These results indicate that, in the context of the ICB regimen, PARP11 inhibition can be used to prevent and treat HPD.

## DISCUSSION

This study demonstrates that upregulation of PARP11 in cancer cells results in increased MARylation, ubiquitination, and degradation of MHC-I, leading to protection of cancer cells from CTLs and accelerated tumor growth. Furthermore, PARP11 activities contribute to tumor cross-resistance to therapies and to the development of HPD. Conversely, pharmacologic inhibition or genetic ablation of PARP11 stabilizes MHC-I, sensitizes cancer cells to killing by CTLs, improves the efficacy of anti-cancer therapies in PDAC models, and prevents HPD in the melanoma model. Along with previous reports on the role of PARP11 in T cells (11, 12), these results provide a foundation for a sustained program to develop potent, selective, and clinically relevant PARP11 inhibitors that can be used as monotherapy in HPD patients and incorporated into PDAC treatment regimens.

Induction of PARP11 in T cells occurs in response to diverse stimuli present in the TME, such as adenosine and prostaglandin E; these GPCR-activating stimuli further act through activation of protein kinase A and CREB transcription factors (12). In addition to adenosine, other GPCR agonists were shown to dampen the extent of tissue inflammation and immunopathology in several mouse models (40). Here we demonstrate that, in addition to adenosine, other GPCR activators, such as epinephrine and GLP1, can also upregulate PARP11 (**Fig 1**), resulting in downregulation of MHC-I and protection of cancer cells from the cytotoxic effects of specific CTLs (**Fig 3, S3**). Adrenergic stress, driven by systemic catecholamine release or by local mediators of the sympathetic nervous system, shapes an immunosuppressive TME and promotes tumor growth and progression (22, 23, 36). The putative contribution of PARP11 induction to these important phenomena is an intriguing possibility that warrants future study.

Given that more than a third of all pharmacological agents approved for patient treatments target GPCRs (24, 25), it is highly plausible that some of these medicines also upregulate PARP11 in cancer cells. We demonstrate that GLP1, which is often over-prescribed or misused in the population without clear indications (26), can upregulate PARP11 and help cancer cells evade CTL control. Although an increased risk of pancreatic cancer in patients using GLP1 was suggested in some (41) but not all (42) reports, additional studies are required to ascertain whether upregulation of PARP11 driven by GLP1, or other GPCR-activating medications may contribute to immunosuppression and tumor progression.

Upregulation of PARP11 in cancer cells promotes tumor growth. Although we cannot rule out the possibility that some cancer cell-intrinsic, non-immune mechanisms also contribute to this phenotype, our results suggest that proteolytic loss of MHC-I and evasion of CTL control play important roles in immunosuppression and tumor growth and progression. It is likely that the latter mechanism, along with previously reported inactivation of CTLs (11) and protection of regulatory T cells (12), form the mechanistic foundation for PARP11-driven immunosuppression in the TME. Whereas we also observed that ITK-7 can decrease tumor myeloid infiltration (**Fig S7A**), the importance of PARP11 in the pro-tumorigenic effects of suppressive myeloid cells, which represent actionable targets for anti-cancer therapy (43), warrants future studies.

Cancer cells engage multiple mechanisms to reduce MHC-I expression and undermine antigen presentation (1), including several that promote MHC-I degradation (31). In PDAC tumors, accelerated lysosomal degradation driven by autophagy receptors (such as NBR1 and p62) represents the major actionable pathway (33). The results presented here characterize PARP11-driven MARylation and ubiquitination of MHC-I as a long-sought regulatory mechanism that recruits autophagy receptors to MHC-I and targets it for lysosomal sorting and accelerated proteolysis. Future studies are required to identify the E3 ligases that facilitate ubiquitination of MHC-I upon its MARylation by PARP11 and to develop inhibitors of these E3 ligases that could be used alongside PARP11 inhibitors to restore MHC-I levels in pancreatic cancers. The feasibility of such approaches is highlighted by our preclinical data demonstrating the efficacy of ITK-7 in restoring MHC-I levels in vivo and overcoming PDAC initial resistance to chemotherapy and acquired resistance to RAS inhibition (**Fig 6**).

Previous studies have demonstrated that PARP11 inactivation increases T cell production of IFN-γ (12), raising concerns about aggravation of HPD. However, our current results suggest that PARP11 activity contributes to the development of HPD. Conversely, PARP11 inactivation not only prevents HPD but can be effective in treating this lethal complication of ICB therapy (**Fig 7**). It is plausible that PARP11 inhibition may reduce NAD+ consumption (19), whose depletion plays a key role in HPD pathogenesis (18). Additional contribution of decreased myeloid infiltration in response to ITK-7 (**Fig S7A**) should also be tested in future studies. Lastly, the development of novel clinically relevant PARP11 inhibitors will help not only to delineate the mechanisms of PARP11-imposed immunosuppression but also to pave the way for the treatment of HPD in melanoma and overcoming chemotherapy, immunotherapy, and targeted therapy resistance in PDAC patients.

## ACKNOWLEDGEMENTS

This work was supported by NIH/NCI grants R01 CA288849 (to S.Y.F. and M.S.C.) and R01 CA288849 (to V.S.S. and S.Y.F.) as well as by the Penn Pancreatic Cancer Research Center. We thank Drs. Mirella L. Meyer-Ficca and Ralph G. Meyer (Utah State University) and Dr. Weiping Zou (University of Michigan) for reagents. We are grateful to Drs. Matthew J. Atherton and Antonella Rotolo and all the members of Fuchs, Guo, Koumenis and Fan labs for critical discussion.

## AUTHOR CONTRIBUTIONS

S.Y.F., M.S.C., V.S.S., Y.S., Y.T. and V.T.S. conceived the study and designed the research. Y.S., Y.T. and V.T.S. performed most of the experiments with the help of A.H., R.B., Q.T.B., J.H.L., R.G., and E.A.C., and interpreted the data. S.Y.F. and Y.S. wrote the manuscript. W.G., J.A.D., Y.F., C.K., T.B., and B.Z.S. helped to design the research, discussed the results, and contributed to the manuscript writing.

## MATERIALS AND METHODS

### Study approvals

All animal experiments were approved by the Institutional Animal Care and Use Committee of the University of Pennsylvania (protocol 803995) and were carried out in accordance with the IACUC guidelines.

### Cell lines

Human PDAC PANC1, MP2 and HPAC cell lines, mouse colon adenocarcinoma MC38 and CT26 cell lines and mouse melanoma B16F10 were purchased from Kerafast and ATCC. The Yumm1.7 cell line was a generous gift from Dr. Weiping Zou (University of Michigan). Mouse pancreatic ductal adenocarcinoma MH6419c5 and MH6499c4 cell lines were previously described (44). Mouse pancreatic ductal adenocarcinoma 4662 cell lines were previously described (45). Human HEK293T cell line was purchased from ATCC.

### Animals

C57BL6^-^ and Balb/c mice were obtained from the Jackson Laboratory. The OT-I mice used to generate OT-I CTLs were obtained from Jackson Labs (C57BL/6-Tg(TcraTcrb)1100Mjb/J, stock # 003831). Rag1-null mice were also from Jackson Lab (B6.129S7-Rag1tm1Mom/J, Stock #002216). C57BL6 littermate *Parp11*^+/+^ (‘WT’) and *Parp11*^-/-^ mice (a generous gift from Drs. Mirella L. Meyer-Ficca and Ralph G. Meyer (Utah State University) were described previously (46). All animal experiments were conducted using male mice between 8 and 10 weeks of age. Mice were housed under specific pathogen-free (SPF) conditions with a 12-hour light/dark cycle and maintained at a temperature of 20–25°C. They had ad libitum access to water and standard laboratory chow (LabDiet 5010 and MF diet; Animal Specialties & Provisions), in accordance with the guidelines of the American Association for Laboratory Animal Science (AALAS). Littermates from different cages were randomly assigned to experimental groups and were housed under identical environmental conditions. Animal health and welfare were routinely monitored by qualified veterinary staff. The genotyping PCR primers were provided in **Supplementary Table 1.**

### Cell culture

Mouse colon adenocarcinoma MC38, MC38-OVA cells, mouse melanoma B16F10 cells, human HEK293T cells, Mouse PDAC MH6419c5, MH6499c4, 4662 cells, human PDAC PANC1, MP2 and HPAC cells were maintained in DMEM (Gibco, 11965-092) supplemented with 10% fetal bovine serum (FBS) (Cytiva, SH30071.03) and 1% penicillin-streptomycin (Cytiva, SV3007901). Mouse colon adenocarcinoma CT26 cells were cultured in RPMI-1640 (Gibco, 22400089) containing 10% FBS and 1% penicillin-streptomycin. Mouse melanoma Yumm1.7 cells were maintained in DMEM: F12 (Invitrogen, 11320033), 10% FBS, 1% MEM non-essential amino acid solution (Gibco, 11440-076) and 1% penicillin-streptomycin. Mouse OT-1 cells were cultured in RPMI-1640 supplemented with 10% FBS, 1% penicillin-streptomycin, 1mM sodium pyruvate (Gibco, 11360070), 55μM 2-mercaptoethanol (Gibco, 21985-023), and 20ng/mL recombinant mouse IL-2 (Biolegend, 575404).

All cells were maintained at 37°C in a humidified incubator with 5% CO_2_ and were routinely confirmed to be free of mycoplasma contamination. For stimulation experiments, cells were treated with adenosine (100 μM) (Sigma, CAS A4036), epinephrine (10 μM) (selleckchem, S2521), or GLP-1 (7–37) (100 nM) (Phoenix Pharmaceuticals, 028-13) for 4 h unless otherwise indicated. For inhibitor studies, cells were treated with ITK-7 (Sigma, SML2669), MG132 (ABMOLE BIOSCIENCE, M1902), bafilomycin A1 (Cayman Chemical, 11038), or cycloheximide (Thermo Scientific Chemicals, 66-81-9) at the indicated concentrations and time points.

### Reagents and antibodies

ITK-7 was purchased from Sigma-Aldrich (SML2669). RMC-6236 was purchased from MedChemExpress (HY-148439). Kolliphor HS15 was obtained from Sigma-Aldrich (42966-1KG). PEG (Carbowax) was purchased from Fisher Scientific (25322-68-3). Gemcitabine (Thermo Scientific Chemicals, 95058-81-4), adenosine (Sigma, CAS A4036), epinephrine (selleckchem, S2521), GLP-1 (7–37) (Phoenix Pharmaceuticals, 028-13), MG132 (ABMOLE BIOSCIENCE, M1902), bafilomycin A1 (Cayman Chemical, 11038), and cycloheximide (Thermo Scientific Chemicals, 66-81-9) were purchased from standard commercial sources. For immune checkpoint blockade experiments, anti–PD-L1 antibody (clone 10F.9G2) was used. For CD8 depletion studies, anti-CD8a antibody (clone 2.43) was used. Rat IgG2b isotype control antibody (clone LTF-2) was used as control where indicated. Antibodies used for immunoblotting, immunoprecipitation, and flow cytometry included antibodies against PARP11, H2-KbDb, H2-Kd, ubiquitin, Poly/Mono-ADP Ribose, NBR1, P62, TAX1BP1, ubiquitin, CD45, CD3, CD8, CD4, NK1.1, CD49b, CD11b, Ly6G, Ly6C, F4/80, IFNγ, Granzyme B, and perforin. Detailed catalog numbers are listed in **KEY RESOURCES TABLE**.

### Plasmids and sgRNA information

Mouse PARP11 overexpression plasmid (pLV-mPARP11) and empty vector control were obtained from Origene (MR204891L3). The catalytic-deficient HY mutant (H197P/Y229S) was generated by site-directed mutagenesis and verified by Sanger sequencing. For CRISPR-mediated gene knockout, sgRNAs targeting mouse *Parp11* were cloned into the lentiCRISPR v2 vector (Addgene, 52961). The SgRNA sequences are NTC: 5ʹ-ATTGAGAATTCGTTTCAAGG-3ʹ, sgParp11-1: 5ʹ-GACGCACGAGTATAATGAAG-3ʹ and sgParp11-2: 5ʹ-GAACCCTGACGTTCCCTACC-3ʹ. Stable knockout cells were established following lentiviral transduction and puromycin selection.

### Lentiviral transduction and generation of stable cell lines

Lentivirus was produced in HEK293T cells using psPAX2 (Addgene, 12260) and pCMV-VSVG (Addgene, 8454) packaging plasmids. For a 100-mm dish, 4 μg transfer plasmid, 1.3 μg psPAX2, and 2.7 μg pCMV-VSVG were transfected using Lipofectamin2000 (Invitrogen, 11668019) reagent according to the manufacturer’s instructions. Viral supernatants were collected 48 h after transfection and concentrated using Lenti-X concentrator (Takara Bio, 631232). Virus pellets were resuspended in complete medium and used to infect target cells in the presence of polybrene. Stable cell lines were selected using puromycin or blasticidin and confirmed by immunoblot analysis.

For MHC-I restoration experiments, cells were co-transfected with pSBbi-pur-H-2Kb (Addgene 111623), β2-microglobulin expression plasmid (Addgene 15883), and pCMV-SB100X transposase plasmid (Addgene 34879) at a ratio of 20:4:1 using Lipofectamine 2000 according to the manufacturer’s instructions. Following transfection and selection, cell surface H-2Kb/Db expression was analyzed by flow cytometry, and cells with comparable MHC-I surface levels were used for subsequent experiments.

### OT-1 cells isolation and in vitro co-culture assay

OT-1 donor mice (C57BL6-Tg (TcraTcrb)1100Mjb/J, RRID: IMSR_JAX:003831) were purchased from The Jackson Laboratory. Splenocytes were isolated and CD8^+^ T cells were purified using magnetic bead-based isolation. Purified OT-1 CD8^+^ T cells were activated using Ovalbumin_257-264_ (Sigma, S7951) peptide stimulation in complete RPMI medium supplemented with recombinant IL-2. For cytotoxicity assays, MC38-OVA, 4662-OVA and B16F10-OVA cells were used as target cells. Tumor cells were pretreated with adenosine, epinephrine, or GLP-1 (7–37) for 4 h before co-culture with activated OT-1 cells. For 7-AAD-based killing assays, effector and target cells were co-cultured at an E:T ratio of 1:1. Tumor cell death was determined by flow cytometric analysis of 7-AAD positivity. For target cells with Luciferase, OT-1 cells were washed and co-cultured with pre-seeded Luciferase-expressing tumor cells in 96-well plates with a total volume of 200 μL at a ratio as indicated. Target cells alone were seeded in parallel at the same density to quantify spontaneous death luciferase expression (measured in relative luminescent units; spontaneous death RLU). Additionally, target cells with water were included as a control for maximal killing (maximal killing RLU). Following coculture, 100 μl of luciferase substrate (Promega, E6120) was added to the remaining supernatant and cells. Luminescence was measured after a 10-minute incubation. Percent cell lysis was calculated using the formula: % lysis = 100 × (spontaneous death RLU - test RLU) / (spontaneous death RLU - maximal killing RLU).

### Cell proliferation assay

Cell proliferation was measured using the CellTiter-Glo Luminescent Cell Viability Assay (Promega, G9241) according to the manufacturer’s instructions. Cells were seeded in 96-well plates and treated as indicated. Luminescence was measured using a microplate reader and normalized to control groups.

### Quantitative real-time PCR

Total RNA was extracted using RNeasy Plus Mini Kit (QIAGEN, 74134). The mRNA was then reverse transcripted to cDNA via reverse transcription using the High-Capacity RNA-to-cDNA Kit (Applied Biosystems, cat 4388950). Quantitative real-time PCR was performed using SYBR Green Master Mix (Applied Biosystems, A25777). Relative gene expression was calculated using the comparative Ct method and normalized to GAPDH. The expression levels were represented as the relative change between groups. The sequences of all oligonucleotide primers can be found in **Supplementary Table 2**.

### Immunoblotting and immunoprecipitation

Cells were lysed in RIPA buffer supplemented with protease and phosphatase inhibitors. Protein concentration was determined using the Bradford assay. Equal amounts of protein were resolved by SDS-PAGE and transferred to PVDF membranes. The following primary antibodies were used: anti-Parp11 (Thermo Scientific, PA5-28545; dilution 1:1,000), anti-H2-KbDB (Biolegend, 114602, 1:100), anti-H2-Kd (Biolegend, 110602, 1:100), anti-Tubulin (Proteintech, 66031-1-Ig, 1:1,000), anti-Actin(Cell Signaling Technology, 4970S, 1:2,000) anti-NBR1 (Thermo Scientific, PA5-106284,1:1,000) anti-P62 (Cell Signaling Technology, 5114S, 1:1,000), anti-TAX1BP1 (Abcam, Ab176572,1:1.000), anti-ubiquitin (Santa Cruz, sc-8017, 1:1,000), anti-Poly/Mono-ADP Ribose (Cell Signaling Technology, 89190S; dilution 1:1,000). Goat Anti-Rabbit IgG Antibody, Peroxidase Conjugated (Sigma, AP132P; dilution 1:5,000) or Goat Anti-Mouse IgG Antibody, Peroxidase Conjugated (Sigma, AP124P; dilution 1:5,000) was used as the appropriate secondary antibody.

For immunoprecipitation assays, cell lysates were incubated with the indicated antibodies overnight at 4°C followed by Protein A/G agarose beads. Immunoprecipitated proteins were analyzed by immunoblotting.

### Migration and invasion assays

Cell migration and invasion were assessed using Transwell chambers with 8-μm pore size inserts (Corning, 354480). For invasion assays, inserts were pre-coated with Matrigel. For migration assays, no Matrigel coating was used. After incubation, migrated or invaded cells were fixed and stained with crystal violet dissolved in 33% acetic acid. The absorbance at OD600 nm was subsequently measured using a microplate reader for quantification.

### Mouse tumor models

Tumor cells were harvested during logarithmic growth phase, washed twice with PBS, and resuspended in HBSS containing 10% Matrigel (Corning, CB40234) prior to subcutaneous injection into the right flank or orthotopic inoculation.

Studies using the orthotopic colon tumor growth, cancer model were carried out as previously described (47). Briefly, after anesthesia, a 1.5cm nick was made in abdominal wall (around) and the cecum was exteriorized and kept moist using PBS. 25μl of MC38 cell suspension (2 x 10^7^/ml) was injected into the cecal wall using 30G needle and the injection site was covered with cotton swab for 3 min to monitor for leakage. Cecum was gently returned to abdominal wall, then abdominal wall and skin were carefully sutured.

Subcutaneous MC38 and CT26 models were established using 5 × 10^5^ cells per mouse. For MH6419c5 pancreatic tumor models, 3 × 10^5^ cells were used for gemcitabine studies and 2 × 10^5^ cells were used for RMC-6236 studies. Hyperprogressive tumor model was established by injecting 5 × 10^5^ Yumm1.7 cells. Tumor volume was measured every 2–3 days using calipers and calculated using the formula: V = (length × width^2^)/2. Once tumors became palpable and reached comparable sizes (75-100 mm^3^), mice were randomly assigned to treatment groups. Mice were euthanized when tumor volume exceeded 1500 mm^3^ or other humane endpoint criteria according to IACUC guidelines.

### ITK-7, gemcitabine, and RMC-6236 *in vivo* treatment

ITK-7 was dissolved in DMSO and administered by intraperitoneal injection. Gemcitabine was dissolved in PBS and administered by intraperitoneal injection at 120 mg/kg twice weekly. RMC-6236 was formulated in DMSO: PEG400: Kolliphor HS15: sterile water at a ratio of 1:2:1:6 and administered by oral gavage at 25 mg/kg once daily.

### CD8 depletion and immune checkpoint blockade

For CD8 depletion experiments, mice were treated with anti-CD8a antibody (clone 2.43, 200 μg per mouse, i.p.). For anti–PD-L1 treatment, mice received anti–PD-L1 antibody (clone 10F.9G2, 200 μg per mouse, i.p.). Rat IgG2b isotype control antibody (clone LTF-2) was used as control.

### Flow cytometry and tumor-infiltrating immune cell analysis

The tumor tissues were cut into small pieces using dissection scissors, followed by incubation in a digestion solution containing 2 mg/ml Collagenase D (Roche, cat 11088882001) for 30 minutes with continuous agitation (200 RPM) at 37°C. Digestion was halted by adding pre-warmed FACS (HBSS with 2% FBS) and passing it through a 100 μm strainer (Corning, 431752) to obtain a single-cell suspension. The cell pellet was resuspended in 1 mL of lysis buffer at room temperature for 5 minutes to remove red blood cells. The cells were then washed twice with FACS. After that, single cells were incubated with the surface, followed by intracellular staining antibodies if required. All conjugated antibodies were diluted to 1:100-1:200 in FACS buffer before use and incubated with cells at 4°C for 30 mins. All antibodies used for flow cytometry are listed in the **KEY RESOURCES TABLE**. Following staining, cells were washed with FACS and subjected to flow cytometry analysis.

### Quantification and statistical analysis

All statistical analyses were performed using GraphPad Prism 9 (GraphPad Prism Software Inc.). Flow cytometry data were analyzed using FlowJo v10. Data are presented as mean ± SEM unless otherwise indicated. For comparisons between two groups, two-tailed unpaired Student’s t test was used. For comparisons among multiple groups, one-way ANOVA or two-way ANOVA was performed. Tumor growth and survival curves were conducted using a repeated-measure two-way ANOVA (mixed model), followed by Tukey’s multiple comparisons test. Mouse survival data were represented using Kaplan-Meier curves. A P value less than 0.05 was considered statistically significant.

### Data availability

All data are available within the article and its supplementary data files or from the corresponding author upon reasonable request.

## Supplementary Information

### LEGENDS TO SUPPLEMENTARY FIGURES

**Figure S1.**
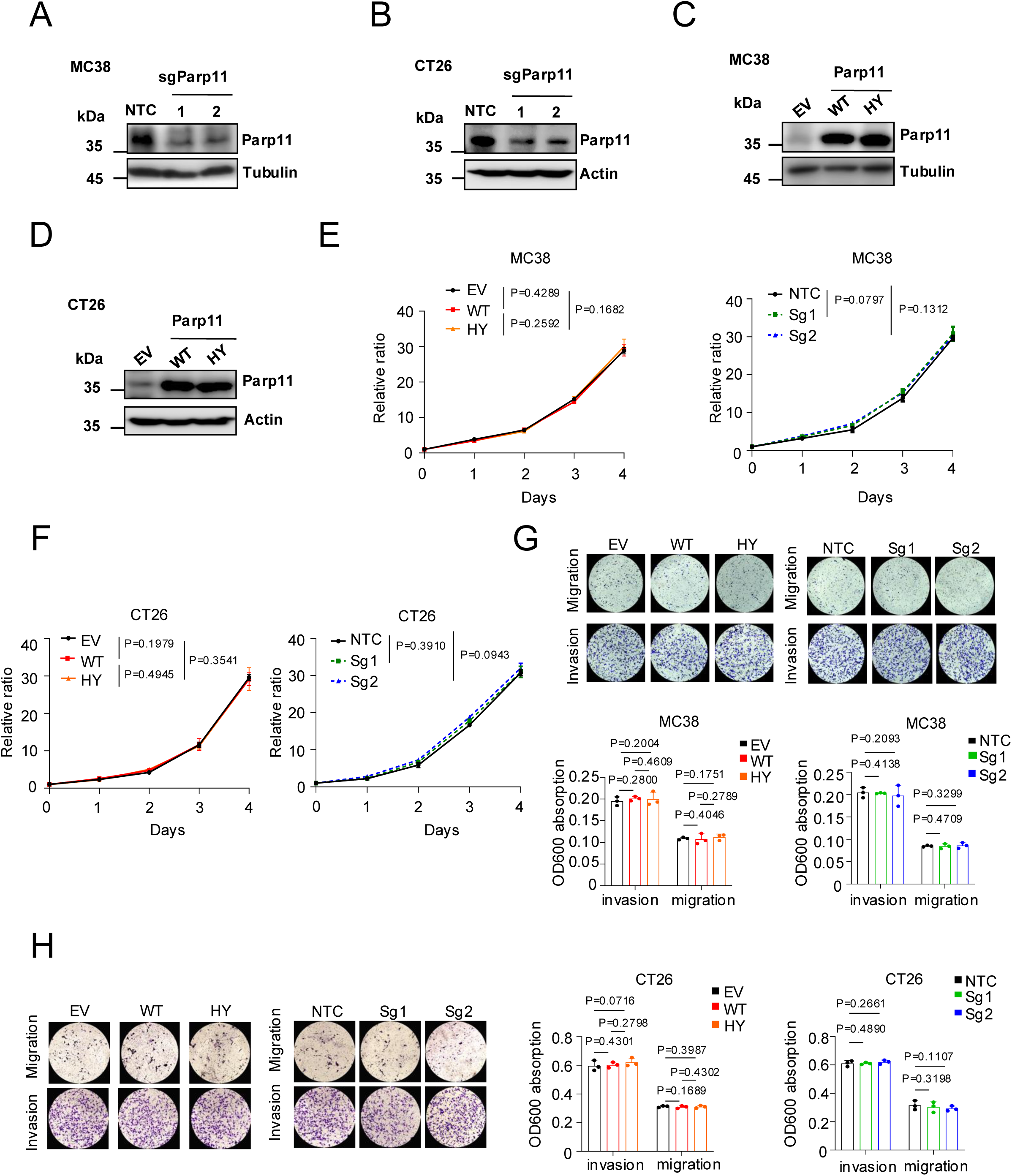
Diverse stimuli in the TME induce PARP11 expression in malignant cells. (A) Immunoblot analysis of PARP11 in MC38 cells expressing two different sgParp11 guides or NTC (Non-Targeting Control); α-Tubulin was used as a loading control. (B) Immunoblot analysis of PARP11 in CT26 cells expressing two different sgParp11 guides or NTC. β-Actin was used as a loading control. (C) Immunoblot analysis of PARP11 in MC38 cells overexpressing wild-type PARP11, catalytic-deficient PARP11 HY mutant (H197P/Y229S), or empty vector (EV) control. α-Tubulin was used as a loading control. (D) Immunoblot analysis of PARP11 in CT26 cells overexpressing wild-type PARP11, PARP11 HY mutant, or EV control. β-Actin was used as a loading control. (E) In vitro proliferation of MC38 cells overexpressing wild-type PARP11, PARP11 HY mutant, or EV control, or MC38 cells expressing sgParp11 or NTC, as described in S1A, C (n = 3 per group). Data is presented as fold change relative to the control groups. (F) In vitro proliferation of CT26 cells overexpressing wild-type PARP11, PARP11 HY mutant, or EV control, or CT26 cells expressing sgParp11 or NTC, as described in S1B, D (n = 3 per group). Data is presented as fold change relative to the control groups. (G) Representative images and quantification of Transwell invasion and migration assays in MC38 cells overexpressing wild-type PARP11, PARP11 HY mutant, or EV control, or MC38 cells expressing sgParp11 or NTC (n = 3 per group). (H) Representative images and quantification of Transwell invasion and migration assays in CT26 cells overexpressing wild-type PARP11, PARP11 HY mutant, or EV control, or CT26 cells expressing sgParp11 or NTC (n = 3 per group). Each dot represents one biological replicate. Data are presented as mean ± SEM. Statistical significance was determined by one-way ANOVA followed by Tukey’s multiple-comparisons test for (E) and (F), and by two-tailed unpaired Student’s t test for (G) and (H).

**Figure S2.**
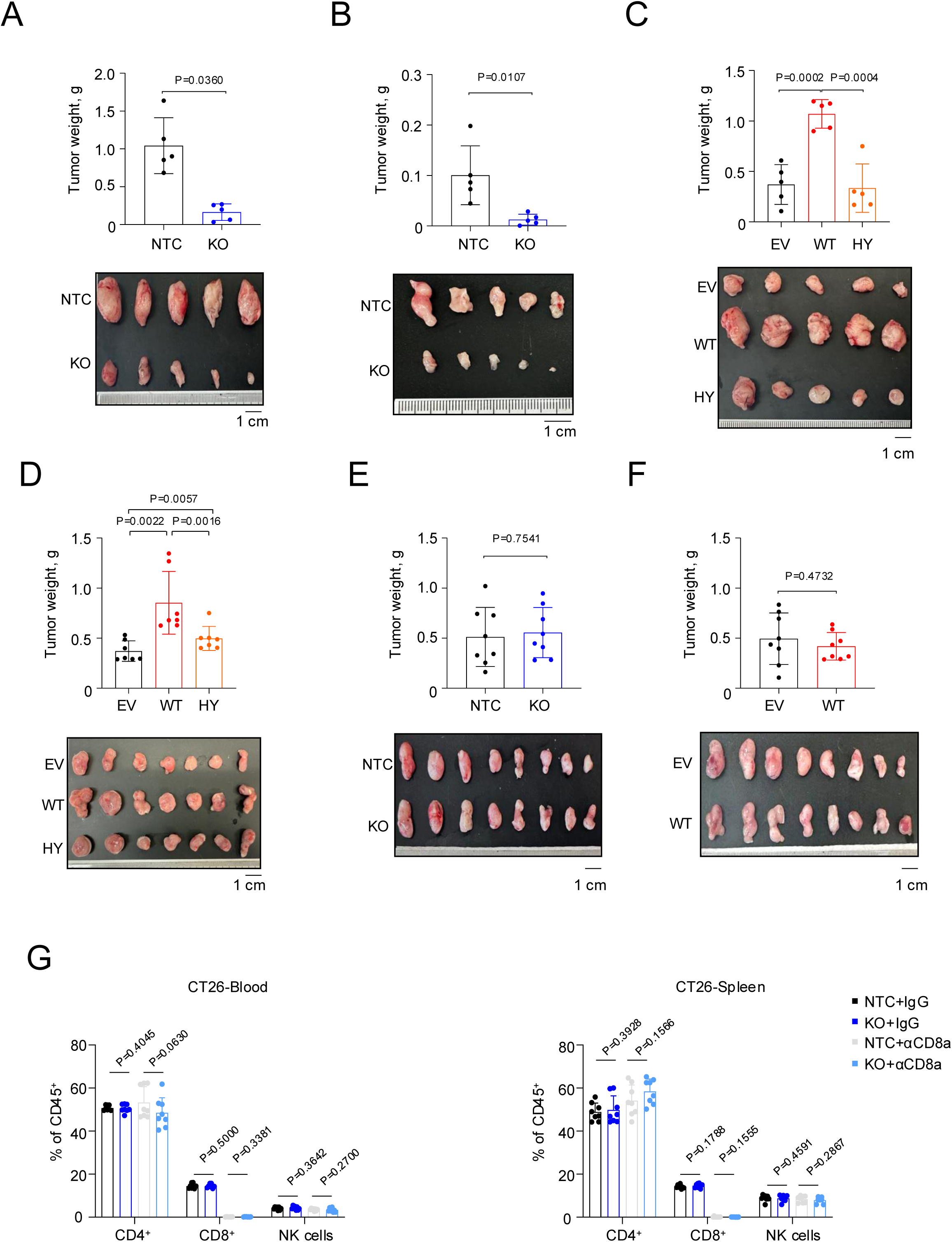
PARP11 promotes tumor growth by evading CD8^+^ T cell-dependent anti-tumor immunity in the TME. (A) Representative tumor images and tumor weights of subcutaneous MC38 tumors expressing sgParp11 or NTC in C57BL6 mice (n = 5 per group), as described in Figure 2A. (B) Representative tumor images and tumor weights of subcutaneous CT26 tumors expressing sgParp11 or NTC in Balb/c mice (n = 5 per group), as described in Figure 2B. (C) Representative tumor images and tumor weights of subcutaneous MC38 tumors overexpressing wild-type PARP11, PARP11 HY mutant, or EV control in C57BL6 mice (n = 5 per group), as described in Figure 2C. (D) Representative tumor images and tumor weights of subcutaneous CT26 tumors overexpressing wild-type PARP11, PARP11 HY mutant, or EV control in Balb/c mice (n = 7 per group), as described in Figure 2D. (E) Representative tumor images and tumor weights of subcutaneous MC38 tumors expressing sgParp11 or NTC in *Rag1*^−/−^ mice (n = 7 per group), as described in Figure 2F. (F) Representative tumor images and tumor weights of subcutaneous MC38 tumors overexpressing wild-type PARP11 or EV control in *Rag1*^−/−^ mice (n = 7 per group), as described in Figure 2G. (G) Quantification of CD4⁺ T cells, CD8⁺ T cells, and NK cells in peripheral blood and spleen from IgG control or anti-CD8α antibody-treated Balb/c mice bearing subcutaneous CT26 tumors expressing sgParp11 or NTC (n = 8 per group), as described in Figure 2H. Each dot represents one biological replicate. Data are presented as mean ± SEM. Statistical significance was determined using two-tailed unpaired Student’s t test (A–G).

**Figure S3.**
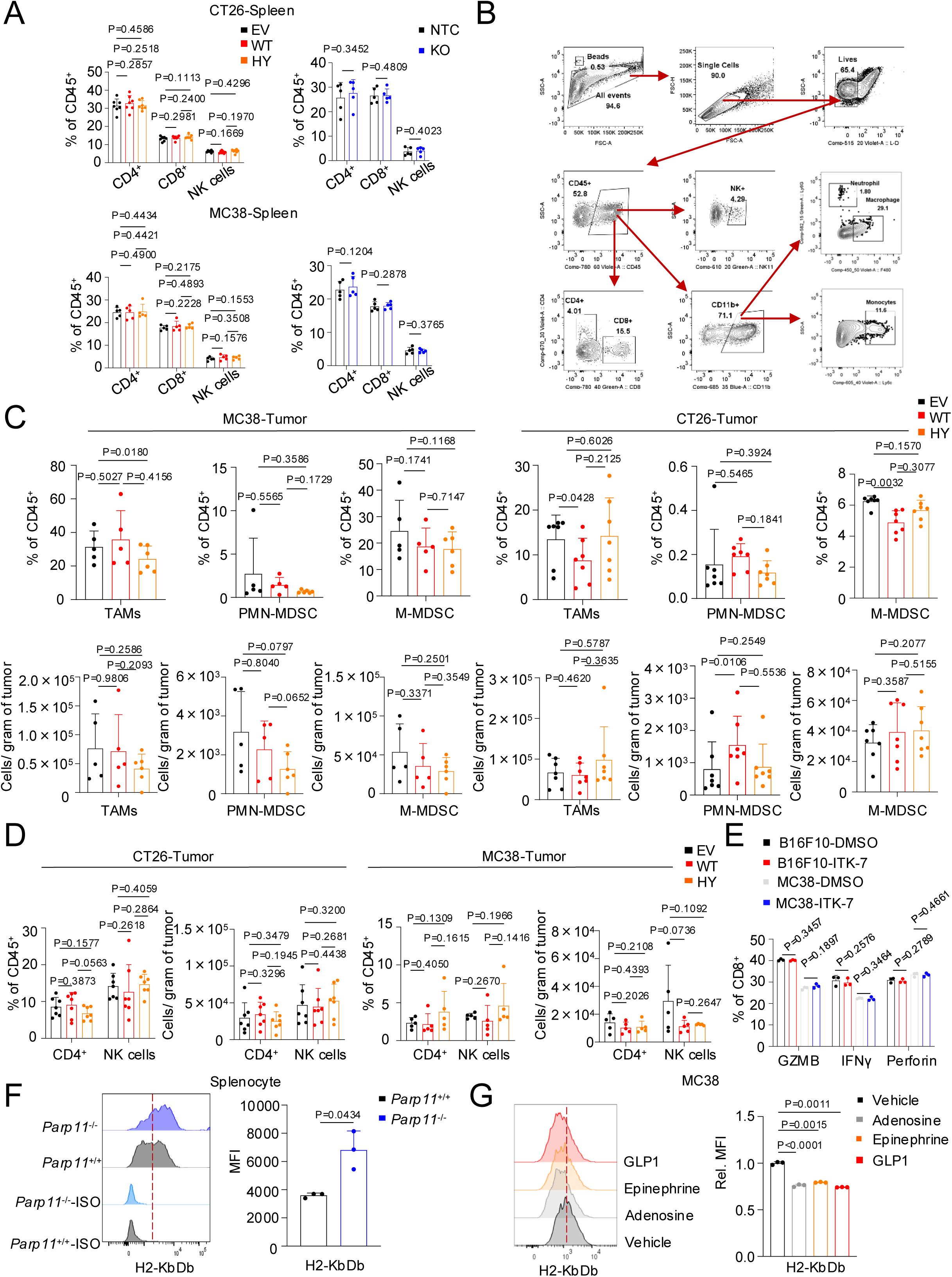
PARP11 suppresses malignant cell surface MHC-I and promotes CD8^+^ T cell-mediated immune evasion. (A) Flow cytometry analysis of the frequency (% of live CD45^+^ cells) of CD4^+^ T cells, CD8^+^ T cells, and NK1.1^+^ or CD49b^+^ cells in the spleens from C57BL6 or Balb/c mice bearing subcutaneous tumors described in Figure 2A–2D. (B) Gating strategy for immune profiling of tumor and splenic tissues. (C) Flow cytometry analysis of the frequency (% of live CD45^+^ cells) and numbers of tumor-associated macrophages (TAMs, CD11b^+^F4/80^+^), polymorphonuclear myeloid-derived suppressor cells (PMN-MDSCs, CD11b^+^Ly6G^+^), and monocytic myeloid-derived suppressor cells (M-MDSCs, CD11b^+^Ly6C^+^), in subcutaneous MC38 or CT26 tumors described in Figure 2C and 2D. (D) Flow cytometry analysis of the frequency and number of CD4^+^ T cells and NK cells in subcutaneous MC38 or CT26 tumors described in Figure 2C and 2D. (E) Flow cytometry analysis of the percentage of IFN-γ^+^, Granzyme B^+^, and perforin^+^ OT-1 T cells in the co-culture system described in Figure 3D (n = 3). (F) Flow cytometry analysis of MFI of H2-KbDb on splenocytes isolated from spleens of WT or *Parp11*^−/−^ mice (n = 3). (G) Flow cytometry analysis of MFI of H2-KbDb on MC38 cells pretreated with adenosine (100 μM), epinephrine (10 μM), GLP-1 (7–37) (100 nM), or vehicle for 4 h (n = 3). Data are presented as fold change relative to the vehicle group. Each dot represents one biological replicate. Data are presented as mean ± SEM. Statistical significance was determined using two-tailed unpaired Student’s t test (A, C–G).

**Figure S4.**
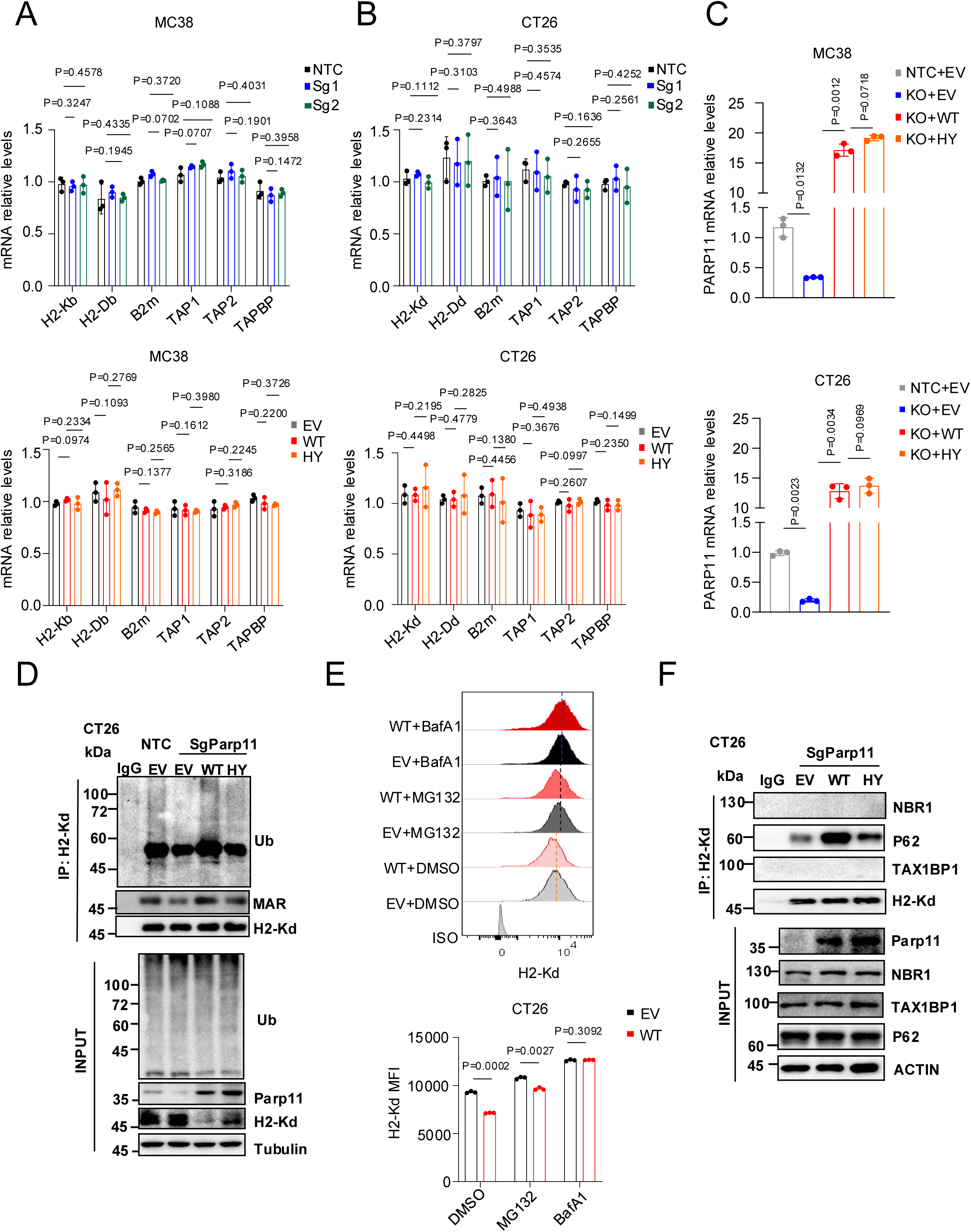
PARP11 promotes MHC-I ubiquitination and lysosomal degradation through MARylation. (A) qRT-PCR analysis of MHC-I, β2-microglobulin, and components of the antigen peptide-presenting machinery in MC38 cells with *Parp11* knockout, wild-type PARP11 overexpression, PARP11 HY mutant overexpression, or corresponding controls, as described in Figure S1A and S1C. Each group contained three biological replicates. Relative mRNA levels were normalized to Gapdh. Data are presented as fold change relative to the control group. (B) qRT-PCR analysis of MHC-I, β2-microglobulin, and components of the antigen peptide-presenting machinery in CT26 cells with *Parp11* knockout, wild-type PARP11 overexpression, PARP11 HY mutant overexpression, or corresponding controls, as described in Figure S1B and S1D. Each group contained three biological replicates. Relative mRNA levels were normalized to Gapdh. Data are presented as fold change relative to the control group. (C) qRT-PCR analysis of Parp11 mRNA expression in MC38 or CT26 cells with endogenous *Parp11* knockout and reconstituted with wild-type PARP11, PARP11 HY mutant, or EV control. Each group contained three biological replicates. Relative mRNA levels were normalized to Gapdh. Data are presented as fold change relative to the control group. (D) CT26 cells with endogenous *Parp11* knockout were reconstituted with wild-type PARP11, PARP11 HY mutant, or EV control were characterized by direct immunoblotting (“Input”, lower panel). Analyses of H2-Kd MARylation and ubiquitination were carried out on MHC-I immunoprecipitates equilibrated for H2-Kd levels (upper panel). (E) Flow cytometry analysis of MFI of H2-Kd on CT26 cells overexpressing wild-type PARP11 or EV control pretreated with MG132 (5 μM), bafilomycin A1 (100 nM), or vehicle for 4 h (n = 3). (F) Immunoprecipitation-immunoblotting analyses of interaction between H2-Kd and NBR1, p62 or TAXBP1 autophagy acceptors in CT26 cells with endogenous *Parp11* knockout were reconstituted with wild-type PARP11, PARP11 HY mutant, or EV control. Direct immunoblotting control (“Input”, lower panel) is also shown. Each dot represents one biological replicate. Data are presented as mean ± SEM. Statistical significance was determined using two-tailed unpaired Student’s t test for (A–C) and (E).

**Figure S5.**
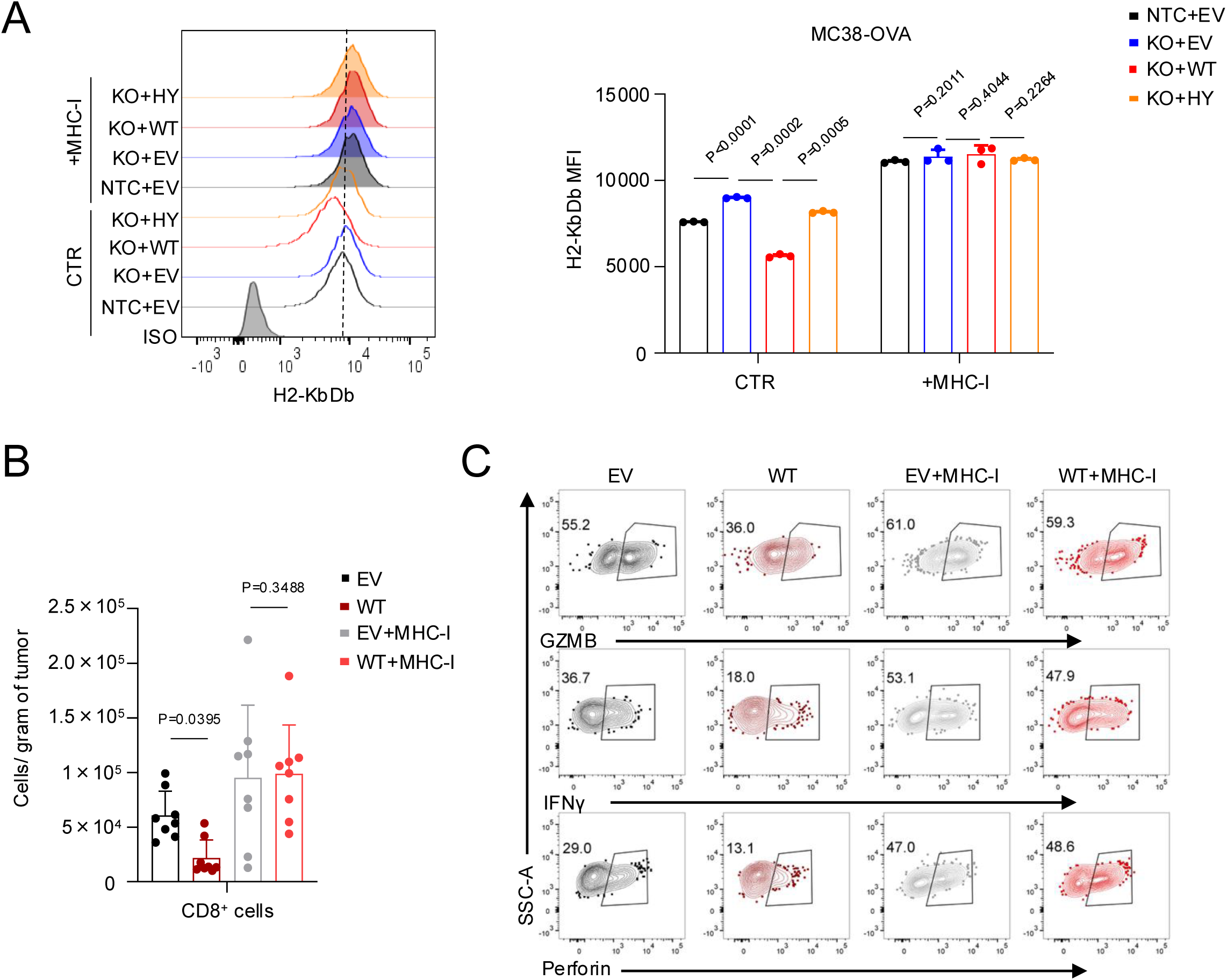
Restoration of MHC-I reverses PARP11-mediated immune evasion. (A) Flow cytometry analysis of MFI of H2-KbDb on MC38-OVA cells treated as described in Figure 5A. (B) Flow cytometry analysis of the number of CD8^+^ T cells per gram of tumor in subcutaneous MC38 tumors treated as described in Figure 5B. (C) Flow cytometry analysis of the percentage of Granzyme B^+^, IFN-γ^+^, and Perforin^+^ CD8^+^ T cells isolated from subcutaneous MC38 tumors in C57BL6 mice treated as described in Figure 5B. Each dot represents one biological replicate. Data are presented as mean ± SEM. Statistical significance was determined using two-tailed unpaired Student’s t test for (A) and (B).

**Figure S6.**
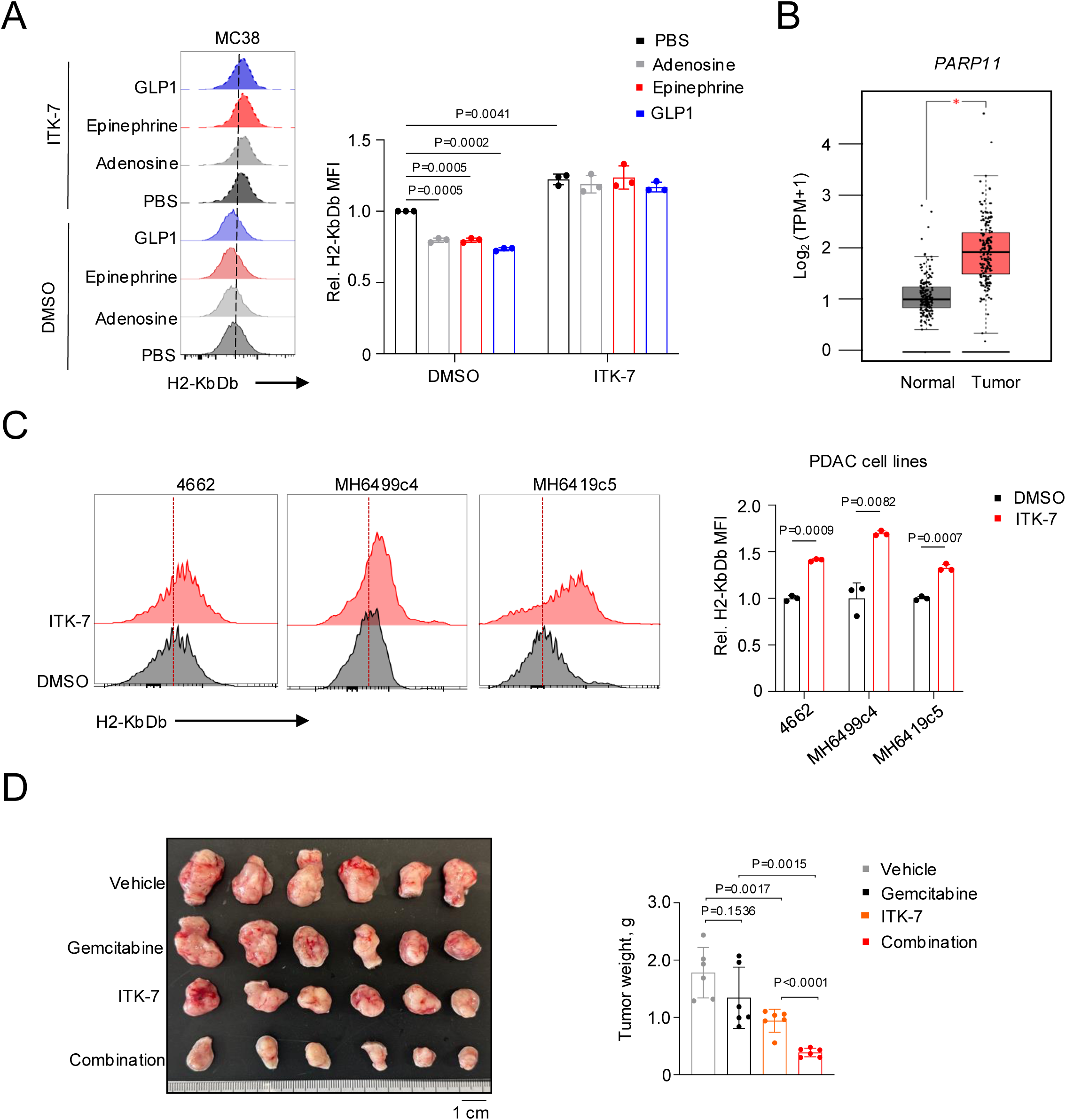
PARP11 inhibition overcomes therapeutic cross-resistance in PDAC tumors. (A) Flow cytometry analysis of MFI of H2-KbDb on MC38 cells. Tumor cells were pretreated with ITK-7 (1 μM, 48 h) or DMSO, followed by adenosine (100 μM), epinephrine (10 μM), or GLP-1 (100 nM), or vehicle for 4 h prior to analysis (n = 3). Data are presented as fold change relative to the vehicle group. (B) Box plot showing PARP11 mRNA expression levels in 179 human pancreatic adenocarcinoma (PAAD) samples and 171 normal tissue samples from TCGA. *P < 0.05 was considered statistically significant. (C) Flow cytometry analysis of MFI of H2-KbDb on 4662, MH6499c4, and MH6419c5 cells pretreated with ITK-7 (1 μM, 48 h) or DMSO. Data are presented as fold change relative to the control group. (D) Representative tumor images and tumor weights of subcutaneous MH6419c5 tumors. Mice were treated as described in Figure 6B. Each dot represents one biological replicate. Data are presented as mean ± SEM. Statistical significance was determined using two-tailed unpaired Student’s t test for (A), (C) and (D).

**Figure S7.**
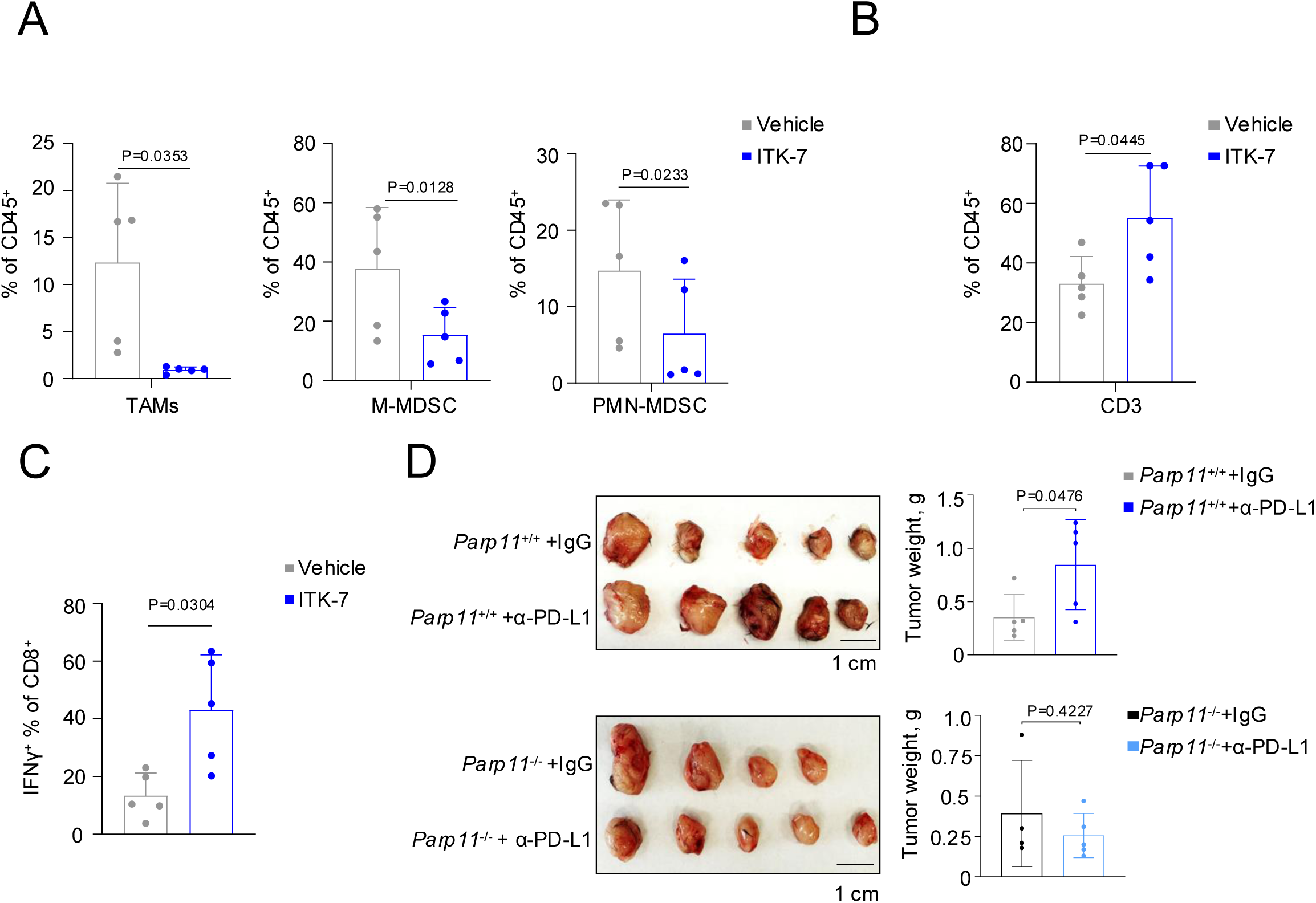
PARP11 inhibition prevents hyper-progressive disease in melanoma. (A) Flow cytometry analysis of frequency of TAMs, PMN-MDSC and M-MDSC cells isolated from subcutaneous YUMM1.7 tumors grown in WT syngeneic mice treated with anti–PD-L1 antibody (200 μg per mouse, intraperitoneally). Mice were concurrently treated with vehicle control or ITK-7 (100 μg per mouse, intraperitoneally, twice a week for 4 weeks; n = 5 per group) as described in 7A. (B) Flow cytometry analysis of the frequency (% of live CD45^+^ cells) of CD3^+^ T cells in subcutaneous YUMM1.7 tumors treated as described in Figure 7A. (C) Flow cytometry analysis of the frequency (% of live CD45^+^ cells) of IFN-γ^+^ CD8^+^ T cells in subcutaneous YUMM1.7 tumors treated as described in Figure 7A. (D) Representative tumor images and tumor weights of subcutaneous YUMM1.7 tumors from mice treated as described in Figure 7B. Each dot represents one biological replicate. Data are presented as mean ± SEM. Statistical significance was determined using two-tailed unpaired Student’s t test for (A–D).

**Supplementary Table 1.**
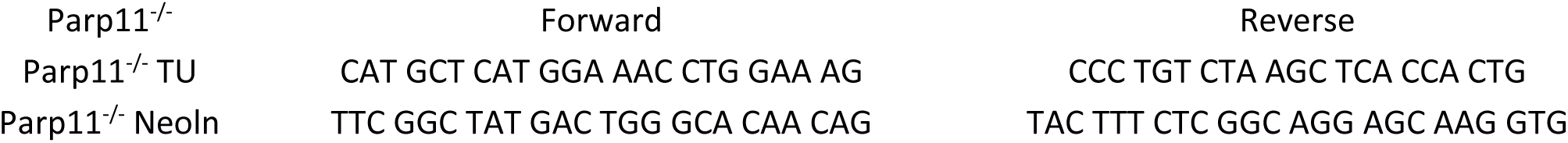
Information of mouse genotying primers.

**Supplementary Table 2.**
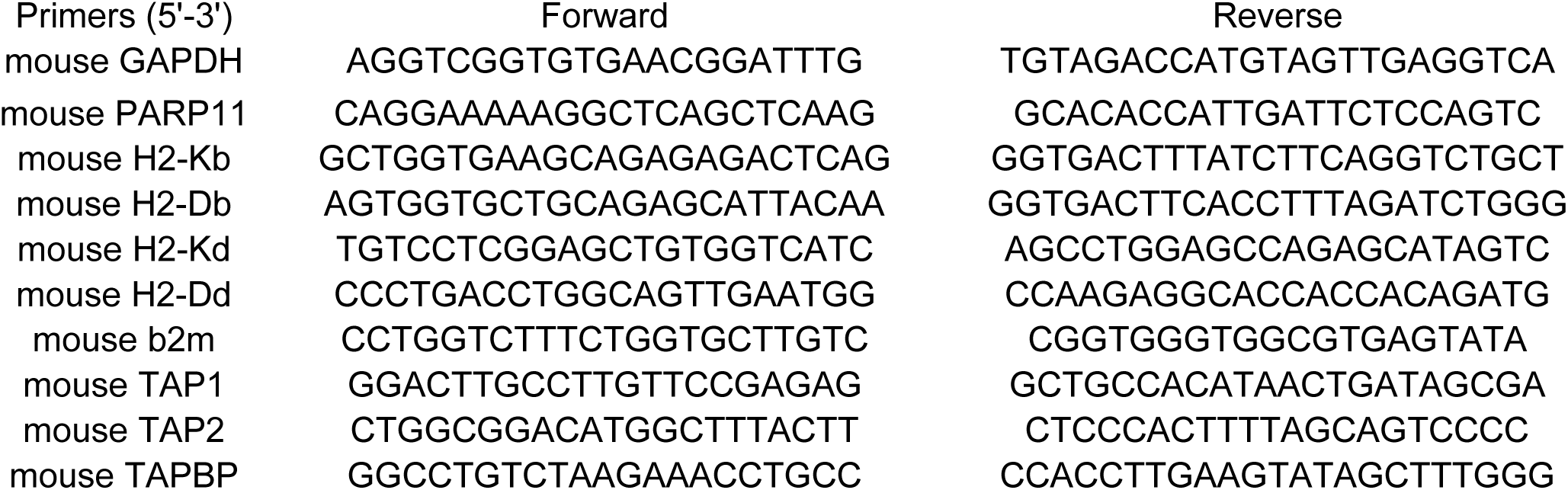
Information of qPCR primers.

